# How Fungi Biosynthesize 3-Nitropropanoic Acid: the Simplest yet Lethal Mycotoxin

**DOI:** 10.1101/2024.02.22.581660

**Authors:** Colin W. Johnson, Masao Ohashi, Yi Tang

## Abstract

We uncovered the biosynthetic pathway of the lethal mycotoxin 3-nitropropanoic acid (3-NPA) from koji mold *Aspergillus oryzae* through genome mining, genetic inactivation and biochemical characterization. The biosynthetic gene cluster (BGC) of 3-NPA, which encodes an amine oxidase and a decarboxylase, is conserved in many fungi used in food processing, although most of the strains have not been reported to produce 3-NPA. Our discovery will lead to efforts that improve the safety profiles of these indispensable microorganisms in making food, alcoholic beverages, and seasoning.

---

Mycotoxins are toxic small secondary metabolites produced by molds (fungi) and can accumulate in food and crops, where they pose health hazards to humans and animals.^[1–3]^ According to USDA, mycotoxins are estimated to impact 25% of the world’s crops and cost US agriculture approximately $1 billion each year.^[4]^ More than 300 mycotoxins have been found to be persistent contaminants of food and agricultural commodities worldwide, including those highlighted by WHO,^[5]^ FDA,^[6]^ and CDC^[7]^ (Figure 1A): trichothecenes, fumonisins and zearalenone from *Fusarium* spp.; citrinin (**2**) from *Penicillium* spp., *Monascus* spp. and *Aspergillus* spp.; aflatoxins (**3**) from *Aspergillus flavus;* ergotamine (**4**) from *Claviceps purpurae;* ochratoxins (**5**) from *Aspergillus* spp. and *Penicillium* spp.; and 3-nitropropanoic acid (3-NPA, **1**) from *Aspergillus oryzae* and other fungi.^[8–10]^ Extensive efforts toward identification and characterization of the biosynthetic gene clusters (BGCs) for the mycotoxins have been undertaken in recent decades.^[11,12]^ The BGCs of many mycotoxins have been successfully identified, as they are all biosynthesized from core enzymes such as polyketide synthases (PKSs), nonribosomal peptide synthetases (NRPSs) and terpene synthases (TSs) (Figure 1A). In contrast, the BGC of the lethal mycotoxin 3-NPA (**1**) has remained elusive.

**Figure 1.**
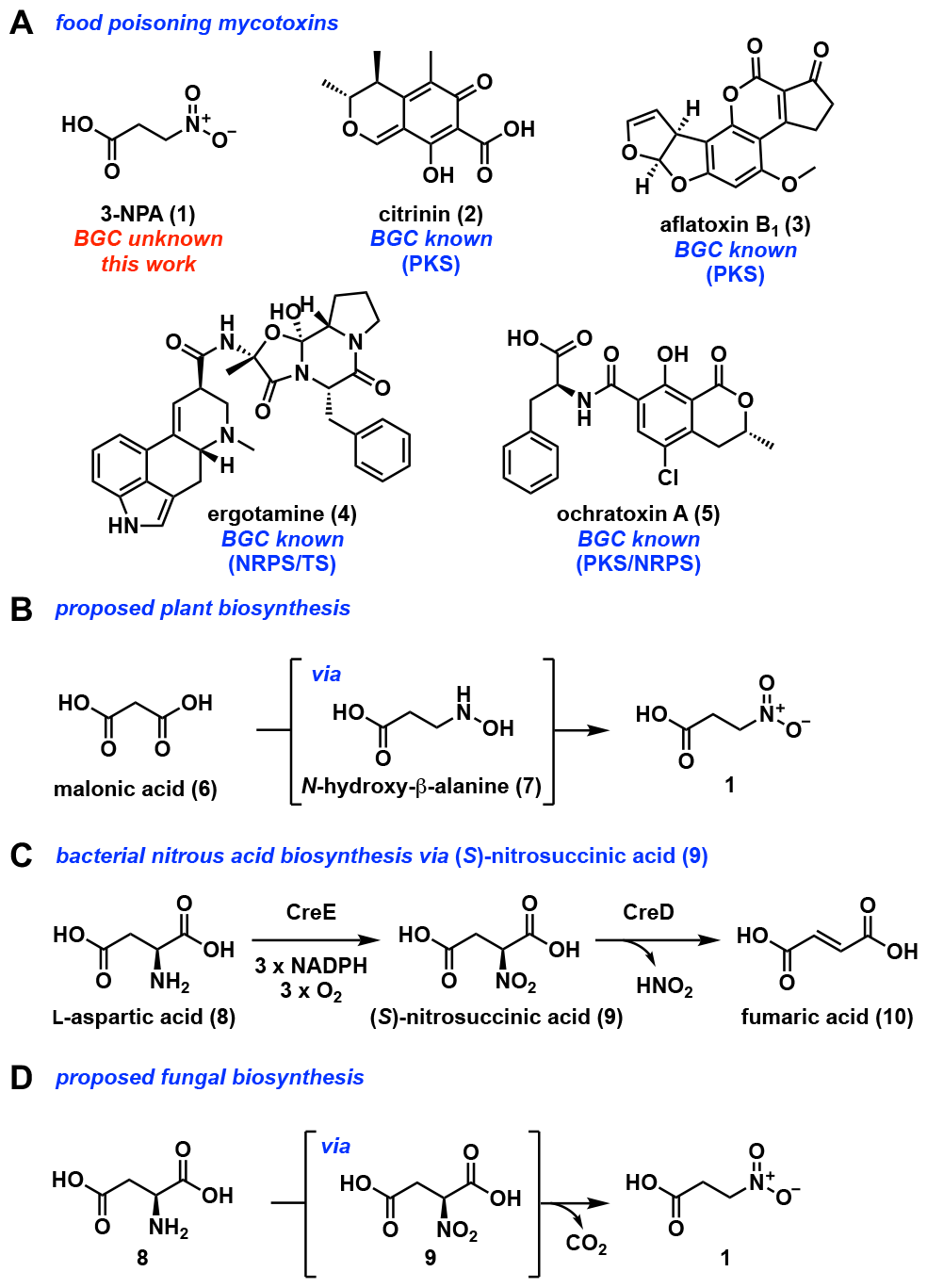
3-NPA is a lethal mycotoxin. (**A**) Mycotoxins with relevance to human health and identified BGC. (**B**) The proposed biosynthesis of **1** in plants starting from malonic acid. (**C**) The bacterial pathway of nitrite production through (*S*)-nitrosuccinic acid. (**D**) The proposed biosynthesis of **1** via (*S*)-nitrosuccinic acid.

3-NPA (**1**) is a deadly neurotoxic nitroalkane found in numerous leguminous plants and fungi.^[13–22]^ **1** is known as an antimetabolite of succinate, and irreversibly inhibits succinate dehydrogenase which disrupts the critical process of mitochondrial oxidative phosphorylation.^[7,23,24]^ The first report of **1** was in 1920,^[25]^ representing the first characterized metabolite to bear the nitro-functionality^[28]^when the structure was later elucidated in 1949.^[26],[27]^ There are significant public health concerns stemming from the ability of industrially relevant molds (*A. flavus*^[18,29,30]^ and *A. oryzae*^[19,21,31]^) used in agriculture, food and enzyme fermentation to produce **1**.^[32]^ In particular, *A. oryzae* is a major organism in the production of fermented food processes^[33]^ including shoyu (fermented wheat and soybeans), miso (rice and soybean), and hamanatto (soybeans). Additionally, modern biotechnology processes including the commercial production of enzymes and flavoring agents such as nucleotides and monosodium glutamate also rely on *A. oryzae* fermentation.^[32]^ Cases of food contaminated with **1** causing human deaths are well documented,^[34]^ with deaths as recent as 2021 by *Arthrinium* spp. (Figure 2A).^[7]^ Therefore, there is urgency to identify genes involved in biosynthesis of **1**.

**Figure 2.**
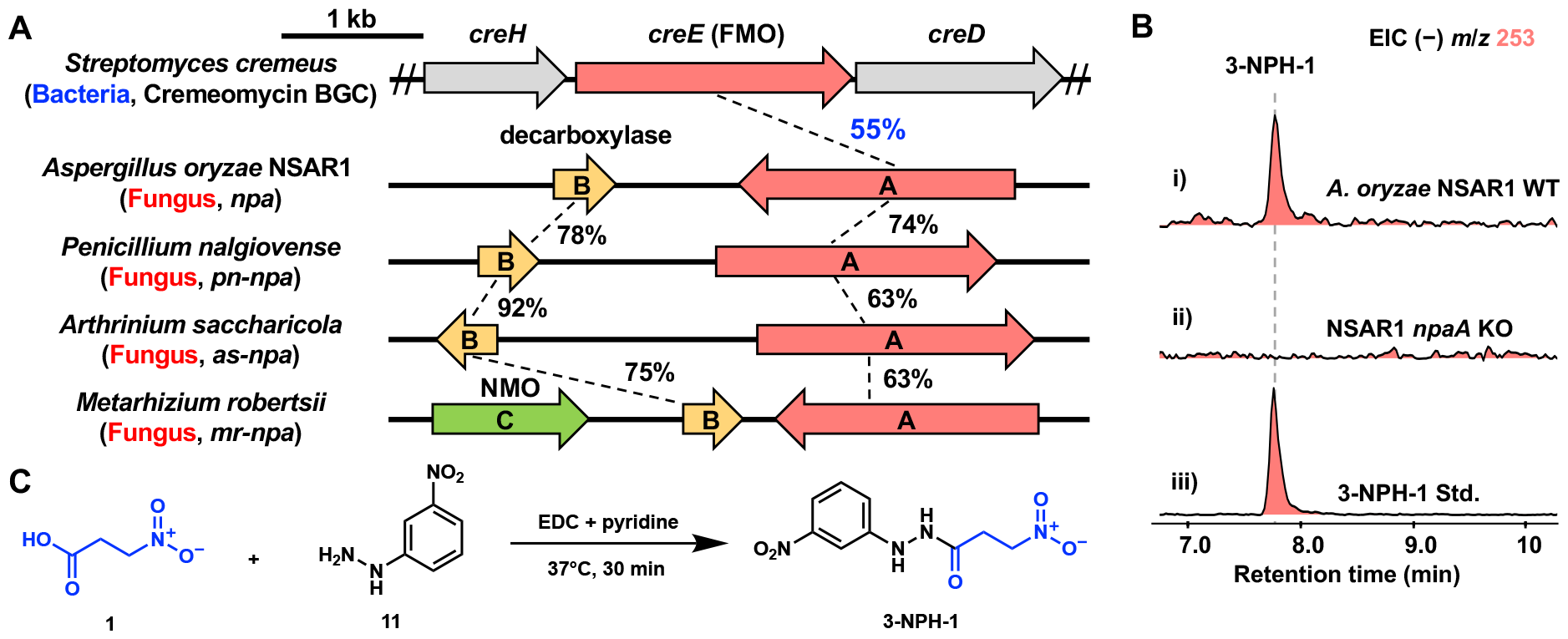
Biosynthesis of **1** by a dedicated biosynthetic gene cluster in fungi. (**A**) Putative BGCs of **1** are not only found in all known producers of **1** such as *A. oryzae* and *A. saccharicola*, but also in fungi not reported to produce **1**, such as *P. nalgiovense* and *M. robertsii* (see Supporting Information). (**B**) LC/MS detection of 3-NPH-**1** from *A. oryzae* WT and abolished detection in the *npaA*-KO strain. (**C**) The synthetic scheme of 3-NPH-**1** using EDC as the condensation reagent.

Isotope-labeling studies of **1** from fungi and plants established the biosynthetic origins are distinct (Figures 1B and 1D).^[20,35–37]^ Whereas the plant pathway starts with malonic acid (**6**) and proceeds via *N*-hydroxy-β-alanine (**7**), fungi initiate the pathway to **1** with L-aspartate (**8**). Baxter *et. al* proposed the iterative oxidation of **8** to generate (*S*)-nitrosuccinic acid (**9**), followed by decarboxylation to **1** (Figure 1D).^[38,39]^ In reexamining this proposal, we noted the oxidation of α-amine of **8** to α-nitro of **9** is identical to the first step of cremeomycin biosynthesis in bacteria, in which a flavin-dependent monooxygenases CreE catalyzes the successive oxidization.^[40]^ **9** then serves as substrate for the lyase CreD, which liberates nitrous acid for N-N bond formation in cremeomycin (Figure 1C). Characterization of FzmM, a close CreE homolog from the fosfazinomycin BGC, led to detection of **1**, which was attributed to the nonenzymatic decarboxylation of **9** to **1**.^[41]^ Based on these prior studies, we hypothesized fungi that produce **1** could encode a CreE-like enzyme possibly acquired through horizontal gene transfer from bacteria.

To test the hypothesis, we searched for a homolog of CreE in the genome of *A. oryzae* and identified a gene encoding a putative flavoenzyme (NpaA) with ∼55% identity to CreE. It is worth noting that CreE/NpaA homologs are widely conserved in not only bacteria, but also fungi (Figure S1). This suggests one or more horizontal gene transfer event(s) of CreE/NpaA homologs between bacteria and fungi might have occurred. Immediately adjacent to *npaA* is a gene encoding a putative decarboxylase (NpaB) annotated as a 4-carboxymucolactone decarboxylase (Figure 2A).^[42]^ The functions of these enzymes are in line with the oxidation-decarboxylation sequence proposed for the biosynthesis of **1** (Figure 1D). Significantly, the juxtaposition of these two genes is not only conserved across all known fungal producers of **1**, but also present in fungi not reported as producers (Figure 2A and Figure S2). Interestingly, species in *Metarhizium* and *Hypoxylon* genera, both not previously reported producers of **1**, contain an additional conserved gene in the cluster, NpaC, which is putatively annotated as a nitronate monooxygenase (NMO) (Figure 2A and Figure S3).^[43]^ Based on this predicted function, NpaC was hypothesized to catalyze the degradation of **1** as a means of resistance, or to metabolize **1** into nitrogen and carbon sources.

The role of *npa* BGC, in particular *npaA*, was first investigated via genetic knockout in *A. oryzae* NSAR1.^[44]^ *A. oryzae* NSAR1 is auxotrophic for nitrate (*niaD*−), adenine (*adeA*−), methionine (*sC*−) and arginine (*argB*−), and is derived from *A. oryzae* RIB40^[45]^ that is often used as a model for strains of *A. oryzae* used in commercial fermentation processes. *A. oryzae* NSAR1 was cultured in Nakamura media for the detection of **1**, based on a previous report of titers as high as ∼1.2 g/L of **1** when *A. oryzae* ATCC12892 was grown in this media.^[19]^ Detection of **1** was achieved through derivatization of the extract with 3-nitrophenylhydrazine (3-NPH) using the reported method to increase MS sensitivity and the retention on the reverse-phase column.^[46]^ Following validation and optimization of this procedure using commercial 3-NPA standard (Figure S5),^[46]^ 3-NPH-**1** (Figures S19-S20 and Table S4) was successfully detected by LC/MS from NSAR1 after 6 days of culturing in Nakamura media, with an approximate titer of ∼4 mg/L (Figure 2B, i, and Figure S10). To directly implicate NpaA in biosynthesis of **1**, a bipartite knockout cassette was then constructed with homology flanking the target site in *npaA*, using *argB* (restoring the Arg auxotrophy) as a selectable marker. Colonies subjected to two sequential passages of selection were then sequenced to genetically verify successful *npaA* knockout via *argB* integration (Figures S6-S8). The production of **1** was completely abolished in the *npaA*Δ::*argB* knockout strain even after prolonged culturing in Nakamura media (> 24 days) (Figure 2B, ii). This suggested that NpaA is indeed the key enzyme to initiate the biosynthesis of **1** as shown in Figure 1D.

NpaA was next expressed and purified from BL21 Star (DE3) (Figure S9). Purified enzyme displayed the characteristic yellow color of flavin-dependent enzymes. Co-factor co-purified with NpaA was identified as FAD by LC/MS analysis of boiled enzyme supernatant (Figure S11). The use of FAD as the co-factor instead of FMN is in agreement with the bacterial homolog CreE.^[40]^ To assay the activity of NpaA, the purified enzyme was mixed with L-aspartate (**8**), and NADPH, followed by derivatization with 3-NPH and LC/MS analysis. As shown in Figure 3A, 3-NPH-**1** can be detected directly from this mixture within 15 min incubation. Alternative amino acid substrates such as L-glutamate, D-aspartate were not accepted by NpaA as a substrate, showing the strict substrate specificity toward L-aspartate. While NADH can be used as a cofactor for reduction of the flavin co-factor, the efficiency is much lower than NADPH as determined by the yield of 3-NPH-**1** (Figure S12). Collectively, these data pinpoint the role of NpaA as (*S*)-nitrosuccinate synthase in the oxidation of **8** to **9**, which can undergo nonenzymatic decarboxylation with the α-nitro group serving as the electron sink.

**Figure 3.**
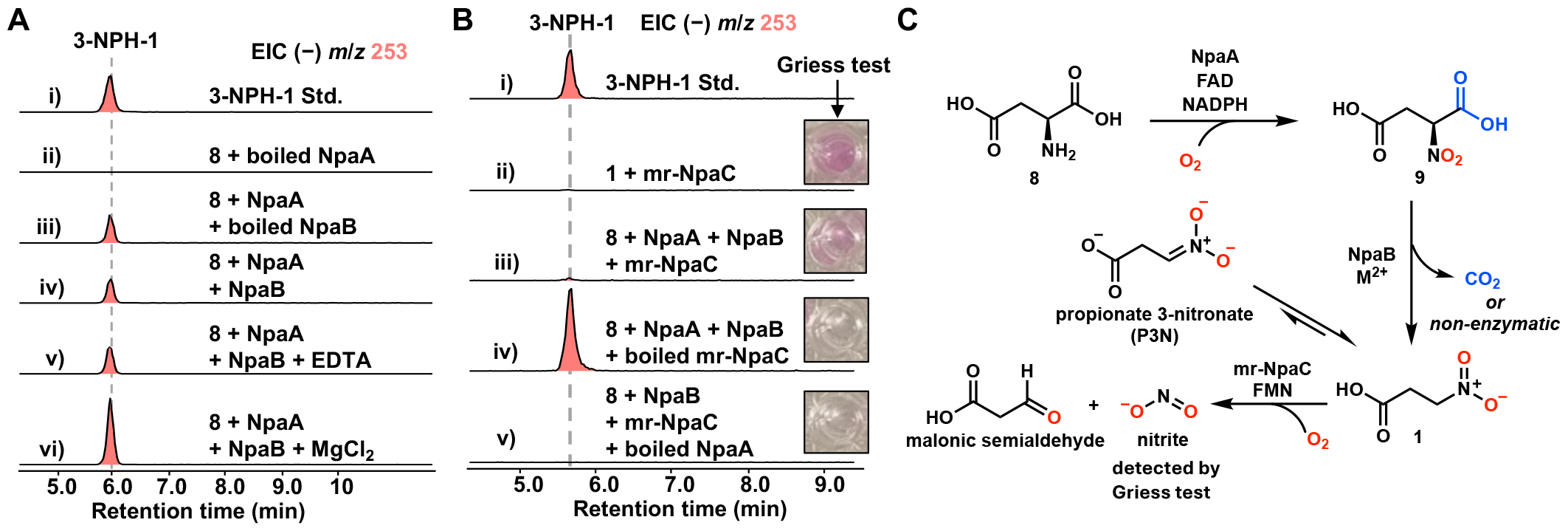
Biochemical characterization of NpaA, NpaB, and mr-NpaC. (A) In vitro characterization of NpaA and NpaB. (B) Detection of nitrite from in vitro reaction of mr-NpaC with **1**. Pinkish color in wells represents the detection of nitrite by Griess reaction. (C) The proposed biosynthetic and degradation pathway of **1**.

While NpaA alone is sufficient to produce **1** under in vitro assay conditions, the putative decarboxylase NpaB colocalized with NpaA could accelerate the decarboxylation of **9** to **1**. NpaB is predicted to be a member of 4-carboxymucolactone decarboxylase family, which consists of the bacterial decarboxylase that catalyzes the cofactor- and oxygen-independent decarboxylation of β-carboxymuconolactone to β-ketoadipate enol lactone in β-ketoadipate pathway.^[42]^ NpaB may facilitate decarboxylation of **9** in a metal-dependent fashion similar to known non-redox decarboxylases.^[47]^ NpaB was expressed and purified from *E. coli* BL21(DE3) (Figure S9). The enzyme was then dialyzed against EDTA-containing buffer to remove any divalent metal ions to become apo-NpaB. A series of combined assays of NpaA and NpaB with and without divalent metal were performed to monitor the production of 3-NPH-**1**. Although the background nonenzymatic decarboxylation of **9** to **1** is also accelerated by divalent metal ions such as Zn^2+^, Mn^2+^ and Mg^2+^ (Figure S14),^[48]^ the production of **1** was the highest when NpaA is paired with NpaB in the presence of MgCl_2_ (Figure 3A, iv-vi, and Figure S13). This notable increase in level of **1** suggests that NpaB is likely the dedicated decarboxylase in the biosynthesis of **1**, using Mg^2+^ as a Lewis acid.

Next, we investigated the role of the conserved, putative nitronate monooxygnease (NMO) in *Metarhizium* (mr-NpaC) and *Hypoxylon* genera (Figure 2A). NMOs are FMN-dependent enzymes that use molecular oxygen to oxidize anionic nitronates or neutral nitroalkanes to the corresponding carbonyl compounds and nitrite (Figure S16).^[43]^ Enzymes capable of catalyzing the oxidation of **1** and/or its anionic form propionate-3-nitronate (P3N) have been discovered from bacteria^[49]^, plants^[50]^ and fungi.^[50–52]^ Studies suggest that NMOs play important roles in not only detoxifing the toxic nitronate, but also allowing the plants and microorganisms to utilize the toxin as source of carbon and nitrogen.^[53]^ Although the sequence identity between mr-NpaC and the characterized fungal NMO from *Neurospora crassa* is moderate (34%) (Figure S17), NpaC may play a similar role in the metabolism of **1**.

To test this possibility, mr-NpaC from *Metarhizium robertsii* was expressed and purified from *E. coli* BL21(DE3) (Figure S9). In agreement with reported fungal NMOs,^[50–52]^ mr-NpaC was co-purified with FMN as judged by LC/MS detection of FMN from denatured mr-NpaC (Figure S15). Upon addition of **1** directly to the reaction mixture containing mr-NpaC, **1** was rapidly consumed as determined through the disappearance of 3-NPH-**1** (Figure 3B, ii and Figure S18), while the consumption was not observed in boiled mr-NpaC control. The formation of nitrite was confirmed by Griess test, as the quenched reaction showed the characteristic pink color following formation of a diazonium salt (Figure 3B, ii). The same conclusions can be drawn from the combined, three-enzyme assay of NpaA, NpaB, and mr-NpaC. Whereas no **1** can be detected in the presence of an active mr-NpaC while the Griess test was positive (Figure 3B, iii), adding denatured mr-NpaC led to accumulation of **1** and a negative Griess test (Figure 3B, iv). As a negative control, adding denatured NpaA led to no detectable **1** as well as a negative Griess test (Figure 3B, v). Collectively, these data confirmed mr-NpaC is indeed a NMO that catalyzes the oxidation of **1** (and P3N) to nitrite and malonic semialdehyde (Figure 3C). While it is possible that mr-NpaC serves as a self-protection mechanism against accumulation of **1** in the producing host, the more likely scenario may be the three enzymes represent an alternative catabolic pathway of aspartate to generate readily metabolizable nitrogen and carbon sources.

Our identification of the biosynthetic pathway of **1** showed minimally only one enzyme, NpaA, is required to make a simple yet deadly mycotoxin. The additional NpaB can accelerate the decarboxylation step and may play a more important role in biosynthesis of **1** in vivo. Prior difficulties in finding the genes for **1** may be due to having the simplest structure among mycotoxins, which makes searching for responsible BGCs using bioinformatics tools difficult. As most of these tools rely on well-studied core enzymes such as PKSs, NRPSs, and TSs, BGCs that do not encode such core enzymes are challenging to find. To leverage knowledge from bacterial pathways in fungal biosynthesis, we used bacterial (*S*)-nitrosuccinate synthase CreE as query to identify the *npa* genes for producing **1** in *A. oryzae*. While such examples of “cross-kingdom genome mining” for secondary metabolism BGCs are still limited, this approach could be fruitful to identify the BGCs of metabolites which are not biosynthesized from canonical core enzymes.^[54]^ One example of this approach was the recent identification of a fungal Pictet-Spenglerase (PS) using the bacterial PS McbB from the marinacarboline biosynthesis.^[55]^

There are significant implications to finally establishing the gene-metabolite link of **1**. In East Asia, mycotoxins produced by koji mold *A. oryzae* is especially concerning since many Asian food and seasonings including soy source and miso are produced from the fermentation of ingredients with *A. oryzae*.^[31,32,45]^ FDA maintains requirement of commercial food additives produced by *A. oryzae* to have levels of mycotoxin, including that of **1**, to be below a threshold.^[32,56]^ BGCs of most mycotoxins produced by *A. oryzae* such as kojic acid,^[57]^ cyclopiazonic acid,^[58]^ and aspirochlorine^[59]^ have been identified, allowing development of modern industrial *A. oryzae* strains with deletions of selected BGCs.^[32]^ In this work, genetic inactivation of *npaA* abolished the production of **1**, and the resulting strain can therefore be considered a safer strain to use for industrial purposes. The *npaA-npaB* pair appears to be widely conserved in many fungi not reported to produce **1** (Figure 2A and Figure S2). Among them, *Penicillium nalgiovense* and *Aspergillus melleus* stand out. *P. nalgiovense* has been used to make mold-fermented meat sausages such as salami, while the traditional production of dried bonito, so-called katsuobushi, involves fermentation using *Aspergillus spp*. such as *A. melleus*.^[60]^ While the detection of **1** in mold-fermented sausages and katuobushi has not been reported to our knowledge, the presence of *npaA-npaB* certainly suggests the fungi are capable of producing **1** (Figure 2A and Figure S2). For such strains, we recommend revisiting the transcriptomes and metabolomes based on results reported here to ensure food safety.

## Data Availability Statement

The data that support the findings of this study are available in the supplementary material of this article.

## Acknowledgments

This work is supported by NIGMS R35GM118056 to YT.

## Notes

### Competing Interest Statement

The authors have declared no competing interest.

## References

[1] J. W. Bennett, M. Klich, Clin. Microbiol. Rev. 2003, 16, 497–516.

[2] J.-D. Bailly, P. Guerre, in Safety of Meat and Processed Meat (Ed.: F. Toldrá), Springer, New York, NY, 2009, pp. 83–124.

[3] A. Moretti, A. F. Logrieco, A. Susca, in Mycotoxigenic Fungi: Methods and Protocols (Eds.: A. Moretti, A. Susca), Springer, New York, NY, 2017, pp. 3–12.

[4] “What are Mycotoxins? : USDA ARS,” can be found under https://www.ars.usda.gov/midwest-area/peoria-il/national-center-for-agricultural-utilization-research/mycotoxin-prevention-and-applied-microbiology-research/docs/what-are-mycotoxins/, (accessed August 2023).

[5] “Mycotoxins,” can be found under https://www.who.int/news-room/fact-sheets/detail/mycotoxins, (accessed August 2023).

[6] “Chemical Hazards,” can be found under https://www.fda.gov/animal-veterinary/biological-chemical-and-physical-contaminants-animal-food/chemical-hazards#Mycotoxins, (accessed August 2023).

[7] T. Birkelund, R. F. Johansen, D. G. Illum, S. E. Dyrskog, J. A. Østergaard, T. M. Falconer, C. Andersen, H. Fridholm, S. Overballe-Petersen, J. S. Jensen, DOI 10.3201/eid2701.202222.

[8] D. Habauzit, P. Lemée, V. Fessard, Food Control 2024, 159, 110273.

[9] R. A. El-Sayed, A. B. Jebur, W. Kang, F. M. El-Demerdash, J. Future Foods 2022, 2, 91–102.

[10] “Mycocentral,” can be found under https://www.mycocentral.eu/, (accessed August 2023).

[11] M. Ferrara, G. Perrone, A. Gallo, Curr. Opin. Food Sci. 2022, 48, 100923.

[12] O. Kolawole, J. Meneely, A. Petchkongkaew, C. Elliott, Fungal Biol. Rev. 2021, 37, 8–26.

[13] R. C. Anderson, W. Majak, M. A. Rassmussen, T. R. Callaway, R. C. Beier, D. J. Nisbet, M. J. Allison, J. Agric. Food Chem. 2005, 53, 2344–2350.

[14] O. Takács, A. Nagyné Nedves, I. Boldizsár, M. Höhn, S. Béni, N. Gampe, Phytochem. Anal. 2022, 33, 1205–1213.

[15] P. Chomcheon, S. Wiyakrutta, N. Sriubolmas, N. Ngamrojanavanich, D. Isarangkul, P. Kittakoop, J. Nat. Prod. 2005, 68, 1103–1105.

[16] J. C. Polonio, M. A. dos S. Ribeiro, S. A. Rhoden, M. H. Sarragiotto, J. L. Azevedo, J. A. Pamphile, Fungal Biol. 2016, 120, 1600–1608.

[17] D.-L. Wei, S.-C. Chang, S.-C. Lin, M.-L. Doong, S.-C. Jong, Curr. Microbiol. 1994, 28, 1–5.

[18] M. T. Bush, Oscar. Touster, Jean. Early. Brockman, J. Biol. Chem. 1951, 188, 685–693.

[19] T. Iwasaki, F. V. Kosikowski, J. Food Sci. 1973, 38, 1162–1165.

[20] J. W. Hylin, H. Matsumoto, Arch. Biochem. Biophys. 1961, 93, 542–545.

[21] A. J. Penel, F. V. Kosikowski, J. Food Prot. 1990, 53, 321–323.

[22] H. Raistrick, A. Stössl, Biochem. J. 1958, 68, 647–653.

[23] T. A. Alston, L. Mela, H. J. Bright, Proc. Natl. Acad. Sci. U. S. A. 1977, 74, 3767–3771.

[24] L. Ming, J. Toxicol. Clin. Toxicol. 1995, 33, 363–367.

[25] Gorter, K. Gaertu Bull Jordin Bot Buitenzorg. 1920, 2, 187–202.

[26] M. P. Morris, C. Pagán, H. E. Warmke, Science 1954, 119, 322–323.

[27] C. L. Carter, W. J. Mcchesney, Nature 1949, 164, 575–576.

[28] R. Parry, S. Nishino, J. Spain, Nat. Prod. Rep. 2010, 28, 152–167.

[29] M. T. Bush, A. Goth, J. Pharmacol. Exp. Ther. 1943, 78, 164–169.

[30] K. C. Marshall, M. Alexander, J. Bacteriol. 1962, 83, 572–578.

[31] S. Deng, K. R. Pomraning, P. Bohutskyi, J. K. Magnuson, Genome Announc. 2018, 6, 10.1128/genomea.00251-18.

[32] J. C. Frisvad, L. L. H. Møller, T. O. Larsen, R. Kumar, J. Arnau, Appl. Microbiol. Biotechnol. 2018, 102, 9481–9515.

[33] C. W. Hesseltine, H. L. Wang, Biotechnol. Bioeng. 1967, 9, 275–288.

[34] B. F. Hamilton, D. H. Gould, D. L. Gustine, in Mitochondrial Inhibitors and Neurodegenerative Disorders (Eds.: P.R. Sanberg, H. Nishino, C.V. Borlongan), Humana Press, Totowa, NJ, 2000, pp. 21–33.

[35] P. D. Shaw, J. A. McCloskey, Biochemistry 1967, 6, 2247–2253.

[36] P. D. Shaw, N. Wang, J. Bacteriol. 1964, 88, 1629–1635.

[37] E. Candlish, L. J. La Croix, A. M. Unrau, Biochemistry 1969, 8, 182–186.

[38] R. L. Baxter, A. B. Hanley, H. W.-S. Chan, S. L. Greenwood, E. M. Abbot, I. J. McFarlane, K. Milne, J. Chem. Soc., Perkin Trans. 1 1992, 2495–2502.

[39] R. L. Baxter, S. L. Smith, J. R. Martin, A. B. Hanley, J. Chem. Soc., Perkin Trans. 1 1994, 2297–2299.

[40] Y. Sugai, Y. Katsuyama, Y. Ohnishi, Nat. Chem. Biol. 2016, 12, 73–75.

[41] Z. Huang, K.-K. A. Wang, W. A. van der Donk, Chem. Sci. 2016, 7, 5219–5223.

[42] D. Parke, Biochim. Biophys. Acta, Protein Struct. 1979, 578, 145–154.

[43] G. Gadda, K. Francis, Arch. Biochem. Biophys. 2010, 493, 53–61.

[44] F. J. Jin, J. Maruyama, P. R. Juvvadi, M. Arioka, K. Kitamoto, FEMS Microbiol. Lett. 2004, 239, 79–85.

[45] M. Machida, K. Asai, M. Sano, T. Tanaka, T. Kumagai, G. Terai, K.-I. Kusumoto, T. Arima, O. Akita, Y. Kashiwagi, K. Abe, K. Gomi, H. Horiuchi, K. Kitamoto, T. Kobayashi, M. Takeuchi, D. W. Denning, J. E. Galagan, W. C. Nierman, J. Yu, D. B. Archer, J. W. Bennett, D. Bhatnagar, T. E. Cleveland, N. D. Fedorova, O. Gotoh, H. Horikawa, A. Hosoyama, M. Ichinomiya, R. Igarashi, K. Iwashita, P. R. Juvvadi, M. Kato, Y. Kato, T. Kin, A. Kokubun, H. Maeda, N. Maeyama, J. Maruyama, H. Nagasaki, T. Nakajima, K. Oda, K. Okada, I. Paulsen, K. Sakamoto, T. Sawano, M. Takahashi, K. Takase, Y. Terabayashi, J. R. Wortman, O. Yamada, Y. Yamagata, H. Anazawa, Y. Hata, Y. Koide, T. Komori, Y. Koyama, T. Minetoki, S. Suharnan, A. Tanaka, K. Isono, S. Kuhara, N. Ogasawara, H. Kikuchi, Nature 2005, 438, 1157–1161.

[46] X. Meng, H. Pang, F. Sun, X. Jin, B. Wang, K. Yao, L. Yao, L. Wang, Z. Hu, Anal. Chem. 2021, 93, 10075–10083.

[47] X. Sheng, F. Himo, Comput. Struct. Biotechnol. J. 2021, 19, 3176–3186.

[48] G. Kubala, A. E. Martell, J. Am. Chem. Soc. 1982, 104, 6602–6609.

[49] S. F. Nishino, K. A. Shin, R. B. Payne, J. C. Spain, Appl. Environ. Microbiol. 2010, 76, 3590–3598.

[50] K. Francis, S. F. Nishino, J. C. Spain, G. Gadda, Arch. Biochem. Biophys. 2012, 521, 84–89.

[51] D. J. Porter, H. J. Bright, J. Biol. Chem. 1987, 262, 14428–14434.

[52] N. Gorlatova, M. Tchorzewski, T. Kurihara, K. Soda, N. Esaki, Appl. Environ. Microbiol. 1998, 64, 1029–1033.

[53] K. Francis, C. Smitherman, S. F. Nishino, J. C. Spain, G. Gadda, IUBMB Life 2013, 65, 759–768.

[54] C. L. M. Gilchrist, H. Li, Y.-H. Chooi, Org. Biomol. Chem. 2018, 16, 1620–1626.

[55] X.-L. Li, Y. Sun, Y. Yin, S. Zhan, C. Wang, Proc. Natl. Acad. Sci. U. S. A. 2023, 120, e2303327120.

[56] “GRAS Notices,” can be found under https://www.cfsanappsexternal.fda.gov/scripts/fdcc/index.cfm?set=GRASNotices&id=1039, (accessed August 2023).

[57] Y. Terabayashi, M. Sano, N. Yamane, J. Marui, K. Tamano, J. Sagara, M. Dohmoto, K. Oda, E. Ohshima, K. Tachibana, Y. Higa, S. Ohashi, H. Koike, M. Machida, Fungal Genet.Biol. 2010, 47, 953–961.

[58] X. Liu, C. T. Walsh, Biochemistry 2009, 48, 8746–8757.

[59] P. Chankhamjon, D. Boettger-Schmidt, K. Scherlach, B. Urbansky, G. Lackner, D. Kalb, H.-M. Dahse, D. Hoffmeister, C. Hertweck, Angew. Chem. Int. Ed. 2014, 53, 13409–13413.

[60] M. J. R. Nout, K. E. Aidoo, in Industrial Applications (Ed.: H.D. Osiewacz), Springer, Berlin, Heidelberg, 2002, pp. 23–47.

