## Supporting Information for "How Fungi Biosynthesize 3-Nitropropanoic Acid: the Simplest yet Lethal Mycotoxin"

#### Table of Contents


**Experimental procedures** **S4**
1. Strains and culture conditions. **S4**
2. General DNA manipulation techniques. **S4**
3. Spectroscopic analyses. **S4**
4. Derivatization of 3-NPA (**1**) with 3-NPH for LC-MS detection. **S5**
5. Synthesis and characterization of **3-NPH-1** standard. **S6**
6. Construction of the *npaA* disruption cassette for knockout in *A. oryzae* NSAR1. **S6**
7. *A. oryzae* NSAR1 protoplast preparation. **S6**
8. Generation of *npaA* knockout in *A. oryzae* NSAR1. **S6**
9. Heterologous expression of *npaA* and *npaB* in *A. nidulans* A1145 ΔSTΔEM. **S7**
10. Expression and purification of NpaA, NpaB, and NpaC from *E. coli*. **S7**
11. *In vitro* reaction of NpaA. **S8**
12. Combo *in vitro* reaction of NpaA and NpaB. **S8**
13. *In vitro* reaction of NpaC. **S8**
14. Time trial *in vitro* reaction of NpaC. **S9**
15. Time trial combo *in vitro* reaction of NpaA, NpaB, and NpaC. **S9**
16. Nitrite detection via Griess Test. **S9**

**Supplementary Tables** **S10**
**Table S1.** Bioinformatic analysis of Npa enzymes. **S10**
**Table S2.** Primers used in this study. **S11**
**Table S3.** Plasmids used in this study. **S12**
**Table S4.** Spectroscopic data of 3-NPH-derivatized 3-NPA standard (**3-NPH-1**). **S13**

**Supplementary Figures** **S14**
**Figure S1.** blastp analysis of NpaA homologs. **S14**
**Figure S2.** Distribution of *npa* BGC in fungi from the NCBI database identified by cblaster. **S16**
**Figure S3.** Example *npa* clusters encoding a third conserved gene *npaC* identified by cblaster. **S21**
**Figure S4.** Multiple sequence alignments of CreE homologs including NpaA and FmzM. **S22**
**Figure S5.** Validation of 3-NPH derivatization for *in vivo* detection of **1** (as **3-NPH-1**). **S23**
**Figure S6.** Scheme of *npaA* disruption cassette design for *npaA* KO in *A. oryzae* NSAR1. **S24**
**Figure S7.** Amplification of *npaA* bipartite knockout fragments. **S25**
**Figure S8.** PCR and sequencing verification of successful *npaA* KO in *A. oryzae* NSAR1. **S26**
**Figure S9.** SDS−PAGE analyses of NpaA, NpaB, and mr-NpaC expressed and purified from *E. coli*. **S27**
**Figure S10.** Standard curve of **3-NPH-1** to estimate the production of **1** in *A. oryzae* NSAR1. **S28**
**Figure S11.** Identification of FAD as cofactor of NpaA. **S29**
**Figure S12.** Identification of NADPH as native co-substrate of NpaA. **S29**
**Figure S13.** Screening EDTA-dialyzed NpaB with divalent metals. **S30**
**Figure S14.** Determination of background non-enzymatic decarboxylation with boiled NpaB. **S31**
**Figure S15.** Identification of FMN as cofactor of mr-NpaC. **S32**
**Figure S16.** Representative example of NMO oxidation yielding carbonyl compounds and nitrite. **S33**
**Figure S17.** Sequence alignment of mr-NpaC with characterized NMO from *Neurospora crassa*. **S34**
**Figure S18.** Time-dependent experiment of 3-NPA consumption and nitrite production by mr-NpaC. **S35**
**Figure S19.** $^{1}H$ NMR spectrum of compound **3-NPH-1** in DMSO-$d_6$. **S36**
**Figure S20.** $^{13}C$ NMR spectrum of compound **3-NPH-1** in DMSO-$d_6$. **S37**

**Sequence Information** **S38**
DNA sequence of *npaA* homolog *as-npaA* gene. **S38**

Protein sequence of NpaA homolog as-NpaA. **S38**
DNA sequence of *npaB* homolog *as-npaB* gene. **S39**
Protein sequence of NpaB homolog as-NpaB. **S39**
DNA sequence of synthesized *mr-npaC* gene. **S39**

**Supplementary References** **S40**

#### Experimental Procedures

##### 1. Strains and culture conditions.

*Aspergillus oryzae* NSAR1 (genotype *niaD*–, *sC*–, *argB*–, *adeA*–)[1] was maintained on potato dextrose agar (PDA, BD) at 28 °C for cell proliferation or in liquid PDB medium (PDA medium without agar) with appropriate supplements (2 g/L ammonium sulphate for *niaD*–, 1.5 g/L methionine for *sC*–, 1.5 g/L arginine for *argB*–, and 0.1 g/L adenine for *adeA*–) for isolation of genomic DNA. The heterologous expression host *Aspergillus nidulans* A1145 ΔSTΔEM was previously developed in our lab.[2] *A. nidulans* A1145 ΔSTΔEM was grown at 28 °C in CD media (10 g/L glucose, 6 g/L $NaNO_3$, 0.52 g/L KCl, 0.52 g/L $MgSO_4{\cdot}7H_2O$, 1.52 g/L $KH_2PO_4$, 1 mL/L trace elements solution, and 20 g/L agar for solid cultivation) with supplementation of 10 mM uridine, 5 mM uracil, 2.5 mg/L riboflavin, and 0.5 mg/L pyridoxine for the collection of spores; or on CD-ST medium (20 g/L starch, 20 g/L casamino acids, 6 g/L $NaNO_3$, 0.52 g/L KCl, 0.52 g/L $MgSO_4{\cdot}7H_2O$, 1.52 g/L $KH_2PO_4$, 1 mL/L trace elements solution, and 20 g/L agar for solid cultivation) for heterologous expression of the gene cluster and RNA extraction. The trace elements solution (100 mL) contained 2.20 g $ZnSO_4{\cdot}7H_2O$, 1.10 g $H_3BO_3$, 0.50 g $MnCl_2{\cdot}4H_2O$, 0.16 g $FeSO_4{\cdot}7H_2O$, 0.16 g $CoCl_2{\cdot}5H_2O$, 0.16 g $CuSO_4{\cdot}5H_2O$, and 0.11 g $(NH_4)_6Mo_7O_{24}{\cdot}4H_2O$. The dropout components for selection for the three expression vectors were uracil/uridine, pyridoxine and riboflavin. *Saccharomyces cerevisiae* JHY686 was used for *in vivo* yeast homologous recombination. *Escherichia coli* TOP10 were used for plasmid propagation. *E. coli* BL21 (DE3) and *E. coli* BL21Star (DE3) were used for protein expression. All *E. coli* strains were cultured in LB (Fisher BioReagents) media (20 g/L agar for solid cultivation) at 37 °C. Carbenicillin or kanamycin was added to the media at the concentration of 100 µg/mL or 50 µg/mL if necessary.

##### 2. General DNA manipulation techniques.

All the DNA manipulations in this study were conducted according to the manufacturer's protocol. *E. coli* TOP10 was used for cloning, following standard recombinant DNA techniques. DNA restriction enzymes were used as recommended by the manufacturer (New England Biolabs, NEB). Genomic DNA from all fungal were extracted by the Quick-DNA™ Fungal/Bacterial Microprep Kit (Zymo Research). PCR for cloning were performed using Q5 High-Fidelity DNA Polymerase (New England Biolabs, NEB). The gene-specific primers are listed in **Table S2**. PCR products were confirmed by DNA sequencing (Primordium Labs). For isolation of RNA from *A. nidulans* transformants, the strains were grown in CD-ST liquid for 4 days at 28 °C. The RNA extraction steps were performed using RiboPure™ Yeast RNA Isolation Kit (Ambion) following the manufacturer's instructions. Residual genomic DNA in the extracts was digested by DNase I (2 U/µL) (Invitrogen) at 37 °C for 4 hours. SuperScript III FirstStrand Synthesis System (Invitrogen) was used for cDNA synthesis with Oligo-dT primers following directions from the user manual.

##### 3. Spectroscopic analyses.

NMR spectra were acquired on a Bruker AV500 spectrometer with a 5 mm dual cryoprobe at the UCLA Molecular Instrumentation Center ($^1$H NMR 500 MHz, $^{13}$C NMR 125 MHz). High resolution mass spectra were acquired on an Agilent 1260 Infinity II LC equipped with an InfinityLab Poroshell 120 EC-$C_{18}$ column (2.7 µm, 3.0 × 50 mm) and a 6545 QTOF high resolution mass spectrometer (UCLA Molecular Instrumentation Center) using the solvent program [1% $CH_3CN{-}H_2O$ 2 min, then 1−95% $CH_3CN{-}H_2O$ (both with 0.1% formic acid, *v/v*) in 9 min followed by 95% $CH_3CN{-}H_2O$ for 6 min at a flow rate of 0.8 mL/min]. LC-MS analyses were also performed on a

Shimadzu 2020 EV LC-MS (Phenomenex Kinetex, 1.7 µm, 2.0 x 100 mm, $C_{18}$ column) using positive and negative mode electrospray ionization with a linear gradient of 5–95% $CH_3CN$-$H_2O$ with 0.1% formic acid (*v/v*) in 15 min followed by 95% $CH_3CN$ for 3 min with a flow rate of 0.3 ml/min.

**4. Derivatization of 3-NPA (1) with 3-NPH for LC-MS detection.**

**General procedure for 3-NPH derivatization:** All reagents for derivatization were freshly prepared prior to use in 3:7 $H_2O$:MeOH as solvent. Briefly, derivatization reactions were performed as follows. 50 µL volumes of sample to be analyzed were sequentially treated with 50 µL of 50 mM 3-nitrophenylhydrazine (3-NPH, Sigma-Aldrich) in 3:7 $H_2O$:MeOH, 50 µL of 50 mM EDC (Oakwood Chemical) in 3:7 $H_2O$:MeOH, and 50 µL of 7% *v/v* pyridine (Acros Organics) in 3:7 $H_2O$:MeOH and mixed thoroughly. Derivatization mixtures were incubated at 37 °C for 30 min and centrifuged at 17,000 *g* for 10 minutes prior to analysis.

**Validation of 3-NPH derivatization strategy:** For 3-NPH derivatization to be a viable method for detection of endogenously produced **1**, control experiments were conducted to verify successful derivatization could be achieved under relevant conditions (**Figure S5**). To ensure **1** could be efficiently extracted and detected from aqueous culture media, a 500 mL flask of Nakamura media (200 mL) was spiked with 5 mg of **1** standard and subjected to identical culture conditions used to culture *A. oryzae* NSAR1 (30 ˚C shaking at 220 rpm). After 6 days, the media was concentrated to a reduced volume (approx. 10 mL) *in vacuo*. 50 µL was removed, derivatized as described above, and analyzed by LC-MS using a Shimadzu 2020 EV LC-MS (Phenomenex Kinetex, 1.7 µm, 2.0 x 100 mm, $C_{18}$ column) using positive and negative mode electrospray ionization with a linear gradient of 5–95% $CH_3CN$-$H_2O$ with 0.1% formic acid (*v/v*) in 15 min followed by 95% $CH_3CN$ for 3 min with a flow rate of 0.3 ml/min.. Recovery and detection of **1** (as **3-NPH-1)** from conditions representative of those that would be used for culturing *A. oryzae* NSAR1 WT concluded validation of the selected 3-NPH derivatization method.

**For detection in WT:** *A. oryzae* NSAR1 (200 mL) was grown in media by Nakamura and Shimoda (1954) (50 g/L sucrose, 20 g/L peptone, 5 g/L $KH_2PO_4$, 2.5 g/L $CaHPO_4$, 2.5 g/L $MgSO_4 \cdot 7H_2O$, pH 6.4)[3] for 6 days and up to 24 days at 30 ˚C shaking at 220 rpm. The cultures were filtered, and the aqueous culture media was concentrated to a reduced volume (approx. 10 mL) *in vacuo* and filtered through 0.22 µM PES filter. 50 µL of the resultant filtered extracts were sequentially treated with 50 µL of 50 mM 3-NPH in 3:7 $H_2O$:MeOH, 50 µL of 50 mM EDC in 3:7 $H_2O$:MeOH, and 7% *v/v* pyridine in 3:7 $H_2O$:MeOH and mixed thoroughly. Derivatization mixtures were incubated at 37 °C for 30 minutes, centrifuged at 17,000 *g* for 10 minutes, and then subjected to LC-MS analysis with a Shimadzu 2020 EV LC-MS (Phenomenex Kinetex, 1.7 µm, 2.0 x 100 mm, $C_{18}$ column) using positive and negative mode electrospray ionization with a linear gradient of 5–95% $CH_3CN$-$H_2O$ with 0.1% formic acid (*v/v*) in 15 min followed by 95% $CH_3CN$ for 3 min with a flow rate of 0.3 ml/min.

**For detection *in vitro*:** 50 µL of 50 mM 3-NPH in 3:7 $H_2O$:MeOH, 50 µL of 50 mM EDC in 3:7 $H_2O$:MeOH, and 7% *v/v* pyridine in 3:7 $H_2O$:MeOH were added sequentially to 50 µL of quenched *in vitro* reaction and mixed thoroughly. Reaction mixtures were incubated at 37 °C for 30 minutes, centrifuged at 17,000 *g* for 10 minutes, and then subjected to LC-MS analysis with an Agilent 1260 Infinity II LC equipped with an InfinityLab Poroshell 120 EC-$C_{18}$ column (2.7 µm, 3.0 × 50 mm) and a 6545 QTOF high resolution mass spectrometer (UCLA

Molecular Instrumentation Center) using the solvent program [1% $CH_3CN-H_2O$ 2 min, then 1−95% $CH_3CN-H_2O$ (both with 0.1% formic acid, *v/v*) in 9 min followed by 95% $CH_3CN-H_2O$ for 6 min at a flow rate of 0.8 mL/min].

**5. Synthesis and characterization of 3-NPH-1 standard.**

A 4-mL dram vial equipped with stir bar was charged with 3-NPA (59.0 mg, 0.50 mmol, 1.0 eq) and diluted 3:7 $H_2O$:MeOH (4.2 mL, 0.12 M) at room temperature. Then added 3-NPH (103 mg, 0.55 mmol, 1.1 eq), EDC (285 mg, 1.50 mmol, 3.0 eq), and pyridine (121 µL, 1.50 mmol, 3.0 eq). Stirred for 16 hours at room temperature. Upon completion by LC-MS, concentrated *in vacuo* and subjected crude residue to HPLC (semi-preparative HPLC was performed on COSMOSIL $5C_{18}$-AR-II (5 µm, f10 x 250 mm, Nacalai Tesque, Inc., Japan) using a flow rate of 4 mL/min, isocratic MeCN–$H_2O$, 25:75) affording title compound as a yellow amorphous solid (22.7 mg, 17.9% yield). HRMS (ESI) calculated $[C_9H_9N_4O_5]^-$ 253.0578, found 253.0594. $^1$H NMR (500 MHz, DMSO) δ 10.12 (s, 1H), 8.47 (s, 1H), 7.53 (ddd, J = 8.0, 2.3, 0.9 Hz, 1H), 7.48 (t, J = 2.2 Hz, 1H), 7.41 (t, J = 8.1 Hz, 1H), 7.12 (dd, J = 8.1, 1.5 Hz, 1H), 5.04 – 4.57 (m, 2H), 3.11 – 2.71 (m, 2H). $^{13}$C NMR (126 MHz, DMSO) δ 169.06, 150.47, 148.65, 130.08, 118.23, 112.83, 105.59, 70.29, 29.74.

**6. Construction of the *npaA* disruption cassette for knockout in *A. oryzae* NSAR1.**

The deletion mutant of NSAR1 Δ*npaA* was constructed using a split-marker approach[4]. Briefly, ~2-kb fragments flanking the targeted deletion region of *npaA* were amplified by PCR (5'-*npaA* with pCJ-30004-F/R and 3'-*npaA* with pCJ-30005-F/R) from *A. oryzae* NSAR1 genomic DNA extract (**Figure S7B**). pTrpC and tTrpC promoter-terminator pair was amplified from in-house plasmid pHyg ArgB was amplified from *A. nidulans* gDNA. The three fragments were joined to create a disruption cassette by using homologous recombination with a linearized XW55 vector in *S. cerevisiae* strain JHY686. The disruption cassette was then split into two by PCR amplification with an overlapping region of ~500 bp spanning *argB* marker (**Figure S6A**).

**7. *A. oryzae* NSAR1 protoplast preparation.**

To prepare protoplasts, spores of *A. oryzae* NSAR1 were inoculated into 50 mL liquid CD media in a 125 mL flask and germinated at 30 ˚C shaking at 220 rpm overnight. The germinated spores were harvested by centrifugation at 3,750 rpm, 4 °C for 20 min and washed two times with 15 mL osmotic buffer (10 mM sodium phosphate buffer, 1.2 M $MgSO_4$, pH 5.8). The mycelia were then resuspended in 10 mL osmotic buffer containing 30 mg of Lysing enzyme from *Trichoderma harzianum* (Sigma-Aldrich) and 20 mg Yatalase (Takara), transferred to a 125 mL flask, and incubated for 6 hrs at 37 °C with gentle shaking at 80 rpm. The cells were then poured into a 50 mL falcon tube and overlaid with 10 mL trapping buffer (0.6 M sorbitol, 0.1 M Tris-HCl, pH 7.0). After centrifugation at 3,750 rpm, 4 °C for 15 min, the protoplasts were obtained in the interface of the two buffers and further washed with double volumes of STC buffer (1.2 M sorbitol, 10 mM $CaCl_2$, 10 mM Tris-HCl, pH 7.5). The protoplasts were then transferred to a sterile 15 mL falcon tube and washed with STC buffer (10 mL, 1.2 M sorbitol, 10 mM $CaCl_2$, 10 mM Tris-HCI pH 7.5). The protoplasts were resuspended in STC buffer (500 µL) and aliquoted for transformation.

**8. Generation of *npaA* knockout in *A. oryzae* NSAR1.**

For knockout of *npaA*, linear PCR-amplified 3'- and 5-*npaA* KO fragments (**Figure S6A**) or empty ArgB-containing plasmid control were added into 60 µL of protoplasts prepared as described above and incubated on ice for 1 hr. Then 600 μL PEG solution [60% PEG4000 (*w/v*), 50 mM $CaCl_2$ and 50 mM Tris-HCl, pH 7.5] was added to the mixture, followed by additional incubation at room temperature for 20 min. Then the mixture was plated on CD-sorbitol medium (CD solid medium with 1.2 M of sorbitol) with appropriate supplements and cultured at 37 °C for 3-5 days until single colonies appeared. Single colonies were subjected to secondary selection after the transformation to ensure genetic purity. The genomic DNA of picked transformants was isolated and checked with diagnostic PCR (**Figure S6B-C**) and the successful integration and disruption *npaA* gene confirmed by DNA sequencing of PCR fragment amplified from outside the target recombination site (**Figure S7A-C**).

**9. Heterologous expression of *npaA* and *npaB* in *A. nidulans* A1145 ΔSTΔEM.**

The general fungal transformation method has been previously described in detail elsewhere.[5] Briefly, to prepare protoplasts spores of *A. nidulans* A1145 ΔSTΔEM were inoculated into 50 mL liquid CD media with appropriate supplements in a 125 mL flask and germinated at 30 ˚C shaking at 220 rpm for ~9 h. The germinated spores were harvested by centrifugation at 3,750 rpm, 4 °C for 20 min and washed two times with 15 mL osmotic buffer (10 mM sodium phosphate buffer, 1.2 M $MgSO_4$, pH 5.8). The mycelia were then resuspended in 10 mL osmotic buffer containing 30 mg of Lysing enzyme from *Trichoderma harzianum* (Sigma-Aldrich) and 20 mg Yatalase (Takara), transferred to a 125 mL flask, and incubated for 6 hrs at 37 °C with gentle shaking at 80 rpm. The cells were then poured into a 50 mL falcon tube and overlaid with 10 mL trapping buffer (0.6 M sorbitol, 0.1 M Tris-HCl, pH 7.0). After centrifugation at 3,750 rpm, 4 °C for 15 min, the protoplasts were obtained in the interface of the two buffers and further washed with double volumes of STC buffer (1.2 M sorbitol, 10 mM $CaCl_2$, 10 mM Tris-HCl, pH 7.5). The protoplasts were then transferred to a sterile 15 mL falcon tube and washed with STC buffer (10 mL, 1.2 M sorbitol, 10 mM $CaCl_2$, 10 mM Tris-HCI pH 7.5). The protoplasts were resuspended in STC buffer (1 mL) and aliquoted for transformation. For each transformation, plasmids were added into 60 μL of protoplasts and incubated on ice for 1 hr. Then 600 μL PEG solution [60% PEG4000 (*w/v*), 50 mM $CaCl_2$ and 50 mM Tris-HCl, pH 7.5] was added to the mixture, followed by additional incubation at room temperature for 20 min. Then the mixture was plated on CD-sorbitol medium (CD solid medium with 1.2 M of sorbitol) and cultured at 37 °C for 2-3 days until single colonies appeared. Isolated transformants were grown in CD-ST media for the 3 days at 28 °C for heterologous expression of the gene cluster and RNA extraction.

**10. Expression and purification of NpaA, NpaB, and NpaC from *E. coli*.**

The *npaA* and *npaB* genes were amplified with overhang primers from the cDNA of the *A. nidulans* construct heterologously overexpressing *npaA* and *npaB*. The *npaC* gene from *Metarhizium robertsii* ARSEF 2575 (*mr-npaC*) without introns was synthesized by IDT Corporation after codon optimization. The PCR product and PCR-linearized pET-28a(+) expression vector were assembled by NEBuilder® HiFi DNA Assembly Master Mix (NEB). The resulting plasmids (**Table S3**) were confirmed by DNA sequencing and transformed into *E. coli* BL21(DE3) cells (for NpaB and NpaC) or *E. coli* BL21Star cells (for NpaA) for $His_6$-tagged protein induction and purification, respectively. The expression and purification of NpaA, NpaB, and NpaC were performed in a similar manner. *E. coli* transformants harboring the corresponding plasmid were grown overnight in LB medium with 50 μg/mL kanamycin at 37 °C, 220 rpm. 5 mL of the overnight seed culture was used to inoculate 1 L of fresh LB medium supplemented with 50 μg/mL kanamycin and incubated at 37 °C until the optical density at 600 nm ($OD_{600}$ value) reached ~0.6. Then culture was submerged in ice water for 25 min, protein expression was induced with 100 μM isopropyl-β-D-thiogalactopyranoside (IPTG) at 15 °C, 220 rpm. After around 20 h, cells were harvested by centrifugation at 5,000 rpm, 4 °C for 15 min, resuspended in 25 mL A10 KPi buffer (50 mM $K_2HPO_4$, 10 mM imidazole, 500 mM NaCl and 5% glycerol, pH 8.0,), and lysed by sonication on ice. The lysate was centrifuged at 14,000 rpm, 4 °C for 30 min to remove the cellular debris. Then the supernatant was subjected to Ni-NTA affinity chromatography at 4 °C for protein purification according to the manufacturer's protocols (GE Healthcare). The purified protein was concentrated and exchanged into storage buffer (50 mM $K_2HPO_4$, 300 mM NaCl, pH 8.0) by using Amicon® Ultra-15 Centrifugal Filters (50 K for NpaA, 10 K for NpaB and 30 K for NpaC). For NpaB, protein was concentrated and exchanged to a 1 mL volume and then subjected to dialysis (12-14 kDa MWCO Spectra/Por™ Dialysis Membrane Tubing, Fisher Scientific) in 500 mL of EDTA dialysis buffer (50 mM $K_2HPO_4$, 300 mM NaCl, 1 mM EDTA, pH 8.0) and gently stirred at 4 °C. Dialysis buffer was exchanged with fresh buffer twice and then allowed to dialyze overnight with slow constant stirring at 4 °C. EDTA was then removed repeating the dialysis procedure described with EDTA-free storage buffer (50 mM $K_2HPO_4$, 300 mM NaCl, pH 8.0). The purity of protein was checked by SDS-PAGE, and the concentration was determined by Bradford method. The purified proteins NpaB and NpaC were flash-frozen in liquid $N_2$ and stored at -80 °C. NpaA rapidly lost activity upon storage and was always used freshly purified as such.

**11. *In vitro* reaction of NpaA.**

**Standard procedure for *in vitro* reaction of NpaA:** 50 μL scale in *vitro* assay of NpaA was performed at 28 °C in 50 mM KPi buffer (pH 8.0), containing 20 μM NpaA, 1 mM L-aspartic acid, and 5 mM NAD(P)H. The reaction was quenched after 15 min or 12 hrs with 50 μL acetonitrile and centrifuged at 17,000 *g* for 10 minutes. The resultant clarified reaction mixture was then subjected to derivatization as described above. LC-MS analysis was performed with an Agilent 1260 Infinity II LC equipped with an InfinityLab Poroshell 120 EC-$C_{18}$ column (2.7 μm, 3.0 × 50 mm) and a 6545 QTOF high resolution mass spectrometer (UCLA Molecular Instrumentation Center) using the solvent program [1% $CH_3CN-H_2O$ 2 min, then 1−95% $CH_3CN-H_2O$ (both with 0.1% formic acid, *v/v*) in 9 min followed by 95% $CH_3CN-H_2O$ for 6 min at a flow rate of 0.8 mL/min].

**12. Combo *in vitro* reaction of NpaA and NpaB**.

**Standard procedure for combo *in vitro* reactions of NpaA and NpaB:** 50 μL scale combo *in vitro* assays of NpaA and EDTA-dialyzed NpaB were performed at 28 °C in 50 mM KPi buffer (pH 8.0), containing 50 μM NpaA, 25 μM NpaB, 1 mM L-aspartic acid, 5 mM NADPH, and 25 μM $MgCl_2$, $MnCl_2$, or $ZnCl_2$ or 1 mM EDTA. The reactions were quenched after 15 mins, 30 mins, or 12 hrs with 50 μL acetonitrile and centrifuged at 17,000 *g* for 10 minutes. The resultant clarified reaction mixture was then subjected to derivatization as described above. LC-MS analysis was performed with an Agilent 1260 Infinity II LC equipped with an InfinityLab Poroshell 120 EC-$C_{18}$ column (2.7 μm, 3.0 × 50 mm) and a 6545 QTOF high resolution mass spectrometer (UCLA Molecular Instrumentation Center) using the same gradient method described above.

**Determining background non-enzymatic decarboxylation:** To determine background non-enzymatic decarboxylation of the unstable NpaA product (nitrosuccinic acid) to **1**, identical reactions to those described above were conducted in tandem with boiled NpaB.

**13. *In vitro* reaction of NpaC.**

100 μL scale *in vitro* assay of NpaC was performed at 28 °C in 50 mM KPi buffer (pH 8.0), containing 20 μM NpaC and 1 mM 3-NPA standard for 12 hrs. The reaction was quenched with 100 μL acetonitrile and centrifuged

at 17,000 *g* for 10 minutes. The resultant clarified reaction mixture was split for both nitrite detection with Griess Test and derivatization as described above for LC-MS analysis. LC-MS analysis was performed with an Agilent 1260 Infinity II LC equipped with an InfinityLab Poroshell 120 EC-$C_{18}$ column (2.7 µm, 3.0 × 50 mm) and a 6545 QTOF high resolution mass spectrometer (UCLA Molecular Instrumentation Center) using the same gradient method described above.

**14. Time trial *in vitro* reaction of NpaC.**

100 µL scale time trial *in vitro* assays of NpaC were performed at 28 °C in 50 mM KPi buffer (pH 8.0), containing 20 µM NpaC and 1 mM 3-NPA standard. The reactions were quenched at 15 min, 30 min, 1 hr, 2 hr, 4 hr, and 12 hr timepoints with 100 µL acetonitrile and centrifuged at 17,000 *g* for 10 minutes. The resultant clarified reaction mixture was split for both nitrite detection with Griess Test and derivatization as described above for LC-MS analysis. LC-MS analysis was performed s with an Agilent 1260 Infinity II LC equipped with an InfinityLab Poroshell 120 EC-$C_{18}$ column (2.7 µm, 3.0 × 50 mm) and a 6545 QTOF high resolution mass spectrometer (UCLA Molecular Instrumentation Center) using the same gradient method described above.

**15. Time trial combo *in vitro* reaction of NpaA, NpaB, and NpaC**.

100 µL scale *in vitro* assays of NpaA, NpaB, and NpaC were performed at 28 °C in 50 mM KPi buffer (pH 8.0), containing 25 µM Npa A, 25 µM EDTA-dialyzed NpaB, 25 µM NpaC, 1 mM L-aspartic acid, 500 µM NADPH, and 25 µM $MgCl_2$, $MnCl_2$, or $ZnCl_2$ or 1 mM EDTA for 1 hr. The reactions were quenched with 100 µL acetonitrile and centrifuged at 17,000 *g* for 10 minutes. The resultant clarified reaction mixture was split for both nitrite detection with Griess Test and derivatization as described above for LC-MS analysis. LC-MS analysis was performed with an Agilent 1260 Infinity II LC equipped with an InfinityLab Poroshell 120 EC-$C_{18}$ column (2.7 µm, 3.0 × 50 mm) and a 6545 QTOF high resolution mass spectrometer (UCLA Molecular Instrumentation Center) using the same gradient method described above.

**16. Nitrite detection via Griess Test.**

The concentration of nitrite in assay mixtures was determined with a nitrite detection kit. Griess Reagent kit was purchased from ThermoFisher Scientific (Cat. G7921) and used according to the protocol of the manufacturer.

#### Supplementary Tables

**Table S1.** Bioinformatic analysis of Npa enzymes.

| Protein (accession) | Size (aa) | Top hit in NCBI blastp database (% identity) | Putative function |
|---|---|---|---|
| NpaA (XP_001825020.1) | 649 | CreE[6] (A0A0K2JL70.1) (53%) | Nitrosuccinic acid synthase |
| NpaB (XP_001825019.1) | 146 | 4-carboxymuconolactone decarboxylase (P20370.2) (40%) | Nitrosuccinic acid decarboxylase |
| mr-NpaC (XP_007823977.1) | 357 | Nitronate monooxygenase (Q01284.1) (34%) | Nitronate monooxygenase |

**NCBI accession number of enzyme in Figure 2A.**

NCBI Accession number of pn-NpaA: OQE91434

NCBI Accession number of pn-NpaB: OQE91377

NCBI Accession number of mr-NpaA: EXU97730

NCBI Accession number of mr-NpaB: EXU97731

**Table S2.** Primers used in this study.

| Primers | Sequence (5'→3') |
|---|---|
| pCJ-30000-F | CATCCCCAGCATCATTACACCTCAGCATTAATTAAATGAAGTACTCCATTCCAGCCAC |
| pCJ-30000-R | GGAGGACATACCCGTAATTTTCTGGGCATTTAAATGTGGCTGAGTCCATTCTCTGTATAC |
| pCJ-30001-F | GTTTAACTTTAAGAAGGAGATATACCATGAAGTACTCCATTCCAGCCAC |
| pCJ-30001-R | TCAGTGGTGGTGGTGGTGGTGCTCGAGAACTCCAACCTCCAGAACAGTAAC |
| pCJ-30002-F | GAACAATAAACCCCACAGAAGGCATTTTTAATTAAATGTCCACTGATCCGAAAATCAACG |
| pCJ-30002-R | GAGACCCAACAACCATGATACCAGGGGATTTAAATCCTTAGTTCTGCTTCAACGAGGTCG |
| pCJ-30003-F | GTTTAACTTTAAGAAGGAGATATACCATGTCCACTGATCCGAAAATCAACG |
| pCJ-30003-R | TCAGTGGTGGTGGTGGTGGTGCTCGAGAAGCCCAGTTTCCTCAGGTC |
| pCJ-30004-F | CAGCGGCCTGGTGCCGCGCGGCAGCATGAAGTACTCCATTCCAGCCAC |
| pCJ-30004-R | TCAGTGGTGGTGGTGGTGGTGCTCGAGTTCAAACTCCAACCTCCAGAACAG |
| pCJ-30005-F | CAGCGGCCTGGTGCCGCGCGGCAGCATGTCCACTGATCCGAAAATCAACG |
| pCJ-30005-R | TCAGTGGTGGTGGTGGTGGTGCTCGAGTCAAAGCCCAGTTTCCTCAGGTC |
| pCJ-30006-F | CATCATCACAGCAGCGGCCTGGTGCCGCGCGGCAGCATGACGGCGCTGTTGCGTAATG |
| pCJ-30006-R | CTCAGTGGTGGTGGTGGTGGTGCTCGAGTCATAAATGTGCGACTACCGAAGTAAGAATGG |
| pCJ-30007-F | GAAATAATTTTGTTTAACTTTAAGAAGGAGATATACCATGACGGCGCTGTTGCGTAATG |
| pCJ-30007-R | GATCTCAGTGGTGGTGGTGGTGGTGCTCGAGTAAATGTGCGACTACCGAAGTAAGAATGG |
| pCJ-30008-F | ttaattaactcgagggtaccgagctcg |
| pCJ-30008-R | gctagcggccgctaaatggatccgagc |
| pCJ-30009-F | tcgacagaagatgatattgaaggagcact |
| pCJ-30009-R | aagaaggattacctctaaacaagtgtacctgt |
| pCJ-30010-F | gctggagctcggatccatttagcggccgctagcGGAACCCACTAGCCCTACTATCTAG |
| pCJ-30010-R | ccaaaaagtgctccttcaatatcatcttctgtcgaGTAGCTGTGCATCTGCGACTAG |
| pCJ-30011-F | gcacaggtacacttgtttagaggtaatccttcttCCTGTTGAGACAACATGGATTCGAG |
| pCJ-30011-R | gtgaattcgagctcggtaccctcgagttaattaaCCTCATTCCAATTGCTACCATGTCC |
| pCJ-30012-F | GGAACCCACTAGCCCTACTATCTAG |
| pCJ-30012-R | gtcacacaaagcattgatgactgg |
| pCJ-30013-F | gtgtcaacgagtccctatacgacac |
| pCJ-30013-R | CCTCATTCCAATTGCTACCATGTCC |
| pCJ-30014-F | ccacttaacgttactgaaatcatc |
| pCJ-30014-R | atgcttgggtagaataggtaagtc |
| pCJ-30015-F | cagattcaatctgacttacctattctacccaagcatatgagtcattcacgaatcatttc |
| pCJ-30015-R | gctgtttgatgatttcagtaacgttaagtggatcattcaatttttcccttattgacaac |
| pCJ-30016-F | CATGCGATAATACCCTCTGGTGC |
| pCJ-30016-R | CGAGCCTCAGGTATCAAGAAGC |

**Table S3.** Plasmids used in this study.

| Plasmids | Vector | Genes |
| --- | --- | --- |
| pCJ30000 | pArgB | *npaA* knockout cassette (ArgB rescue) |
| pCJ30001 | pYTU | *npaA* (nitrosuccinate synthase) |
| pCJ30002 | pYTP | *npaB* (CMD-like decarboxylase) |
| pCJ30003 | pET-28a(+) | *npaA* (nitrosuccinate synthase) |
| pCJ30004 | pET-28a(+) | *npaB* (CMD-like decarboxylase) |
| pCJ30005 | pET-28a(+) | *mr-npaC* (NMO) |

**Table S4.** Spectroscopic data of 3-NPH-derivatized 3-NPA standard (**3-NPH-1**).

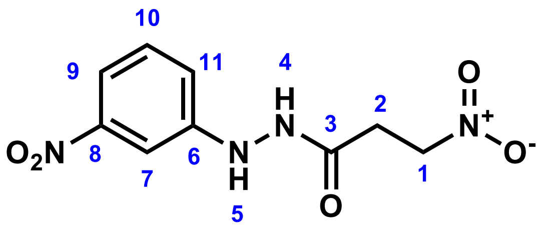


| **3-NPH-1** in DMSO-$d_6$ | | |
|---|---|---|
| **Position** | **$^{13}$C** | **$^{1}$H** |
| 1 | 70.29 | 5.04 – 4.57 (m, 2H) |
| 2 | 29.74 | 3.11 – 2.71 (m, 2H) |
| 3 | 169.06 | - |
| 4 | - | 10.12 (s, 1H) |
| 5 | - | 8.47 (s, 1H) |
| 6 | 150.47 | - |
| 7 | 105.59 | 7.12 (dd, J = 8.1, 1.5 Hz, 1H) |
| 8 | 148.65 | - |
| 9 | 112.83 | 7.53 (ddd, J = 8.0, 2.3, 0.9 Hz, 1H) |
| 10 | 130.08 | 7.41 (t, J = 8.1 Hz, 1H) |
| 11 | 118.23 | 7.48 (t, J = 2.2 Hz, 1H) |

HRMS (ESI) calculated $[C_9H_9N_4O_5]^-$ 253.0578, found 253.0594. $^{1}$H NMR (500 MHz, DMSO) δ 10.12 (s, 1H), 8.47 (s, 1H), 7.53 (ddd, J = 8.0, 2.3, 0.9 Hz, 1H), 7.48 (t, J = 2.2 Hz, 1H), 7.41 (t, J = 8.1 Hz, 1H), 7.12 (dd, J = 8.1, 1.5 Hz, 1H), 5.04 – 4.57 (m, 2H), 3.11 – 2.71 (m, 2H). $^{13}$C NMR (126 MHz, DMSO) δ 169.06, 150.47, 148.65, 130.08, 118.23, 112.83, 105.59, 70.29, 29.74.

### Supplementary Figures

| | | | | | | | | |
|---|---|---|---|---|---|---|---|---|
| hypothetical protein N7449_010936 [Penicillium cf. viridicatum] | Penicillium cf. viridicatum | 947 | 947 | 97% | 0.0 | 73.10% | 644 | KAJ5187942.1 |
| hypothetical protein N7499_013315 [Penicillium canescens] | Penicillium canescens | 947 | 947 | 98% | 0.0 | 72.61% | 644 | KAJ6064635.1 |
| hypothetical protein N7524_004548 [Penicillium chrysogenum] | Penicillium chrysogenum | 947 | 947 | 96% | 0.0 | 73.81% | 644 | KAJ5278395.1 |
| uncharacterized protein N7446_000034 [Penicillium canescens] | Penicillium canescens | 947 | 947 | 98% | 0.0 | 72.61% | 644 | XP_058377780.1 |
| hypothetical protein N7522_002064 [Penicillium canescens] | Penicillium canescens | 947 | 947 | 98% | 0.0 | 72.46% | 644 | KAJ6011709.1 |
| hypothetical protein PENNAL_c0009G03558 [Penicillium nalgiovense] | Penicillium nalgiovense | 947 | 947 | 97% | 0.0 | 73.58% | 644 | OQE91434.1 |
| hypothetical protein N7527_001231 [Penicillium freii] | Penicillium freii | 945 | 945 | 97% | 0.0 | 72.63% | 635 | KAJ5534977.1 |
| hypothetical protein PENPOL_c004G02317 [Penicillium polonicum] | Penicillium polonicum | 944 | 944 | 97% | 0.0 | 72.78% | 635 | OQD66804.1 |
| hypothetical protein N7465_008273 [Penicillium sp. CMV-2018d] | Penicillium sp. CMV-2018d | 944 | 944 | 97% | 0.0 | 72.78% | 635 | KAJ5406989.1 |
| uncharacterized protein N7509_000058 [Penicillium cosmopolitanum] | Penicillium cosmopolitanum | 944 | 944 | 96% | 0.0 | 73.81% | 644 | XP_056494806.1 |
| unnamed protein product [Penicillium nalgiovense] | Penicillium nalgiovense | 944 | 944 | 97% | 0.0 | 73.26% | 644 | CAG8242837.1 |
| FAD-NAD(P)-binding-domain-containing protein [Hypoxylon sp. FL1857] | Hypoxylon sp. FL1857 | 944 | 944 | 94% | 0.0 | 74.27% | 658 | KAI1412133.1 |
| uncharacterized protein N7518_004309 [Penicillium psychrosexuale] | Penicillium psychrosexuale | 943 | 943 | 97% | 0.0 | 72.78% | 644 | XP_057042006.1 |
| hypothetical protein ACN42_g10571 [Penicillium freii] | Penicillium freii | 941 | 941 | 97% | 0.0 | 72.47% | 635 | KUM56632.1 |
| FAD-NAD(P)-binding-domain-containing protein [Aspergillus avenaceus] | Aspergillus avenaceus | 940 | 940 | 100% | 0.0 | 70.45% | 657 | KAE8149226.1 |
| hypothetical protein PENARI_c007G05347 [Penicillium arizonense] | Penicillium arizonense | 939 | 939 | 97% | 0.0 | 72.94% | 644 | XP_022489244.1 |
| hypothetical protein N7530_010544 [Penicillium desertorum] | Penicillium desertorum | 937 | 937 | 95% | 0.0 | 74.24% | 632 | KAJ5462339.1 |
| hypothetical protein PENANT_c026G06017 [Penicillium antarcticum] | Penicillium antarcticum | 937 | 937 | 97% | 0.0 | 72.04% | 644 | OQD81608.1 |
| hypothetical protein CBS147318_3855 [Penicillium roqueforti] | Penicillium roqueforti | 937 | 937 | 97% | 0.0 | 72.47% | 644 | KAI2718745.1 |
| uncharacterized protein N7508_000009 [Penicillium antarcticum] | Penicillium antarcticum | 934 | 934 | 98% | 0.0 | 71.34% | 644 | XP_058324316.1 |
| uncharacterized protein LCP9604111_2521 [Penicillium roqueforti] | Penicillium roqueforti | 934 | 934 | 97% | 0.0 | 72.31% | 644 | XP_038931273.1 |
| hypothetical protein CBS147332_7747 [Penicillium roqueforti] | Penicillium roqueforti | 932 | 932 | 97% | 0.0 | 72.15% | 644 | KAI2701971.1 |
| FAD/NAD(P)-binding protein [Streptomyces sp. NBC_00554] | Streptomyces sp. NBC_00554 | 932 | 932 | 98% | 0.0 | 72.77% | 639 | WUC52579.1 |
| FAD/NAD(P)-binding protein [Streptomyces sp. NBC_00827] | Streptomyces sp. NBC_00827 | 932 | 932 | 98% | 0.0 | 72.61% | 639 | WTB41487.1 |
| FAD/NAD(P)-binding protein [Streptomyces sp. NBC_00564] | Streptomyces sp. NBC_00564 | 930 | 930 | 98% | 0.0 | 72.61% | 639 | WUC10898.1 |
| FAD/NAD(P)-binding protein [Streptomyces sp. NBC_00620] | Streptomyces sp. NBC_00620 | 927 | 927 | 98% | 0.0 | 72.46% | 639 | WP_266463391.1 |
| FAD/NAD(P)-binding protein [Streptomyces sp. NBC_01356] | Streptomyces sp. NBC_01356 | 921 | 921 | 98% | 0.0 | 71.72% | 647 | WP_329547898.1 |
| hypothetical protein ED733_002198 [Metarhizium rileyi] | Metarhizium rileyi | 920 | 920 | 94% | 0.0 | 72.26% | 665 | TWU70941.1 |
| unnamed protein product [Penicillium roqueforti FM164] | Penicillium roqueforti FM164 | 919 | 919 | 97% | 0.0 | 71.68% | 639 | CDM34159.1 |
| uncharacterized protein MAC_09733 [Metarhizium acridum CQMa 102] | Metarhizium acridum CQMa 102 | 919 | 919 | 91% | 0.0 | 73.55% | 663 | XP_007816073.1 |
| FAD-binding protein [Streptomyces sp. Tue6028] | Streptomyces sp. Tue6028 | 919 | 919 | 98% | 0.0 | 70.70% | 670 | PBC64132.1 |
| hypothetical protein J3458_014278 [Metarhizium acridum] | Metarhizium acridum | 919 | 919 | 91% | 0.0 | 73.55% | 663 | KAG8412567.1 |
| FAD/NAD(P)-binding protein [Streptomyces sp. Tue6028] | Streptomyces sp. Tue6028 | 919 | 919 | 98% | 0.0 | 70.70% | 660 | WP_306453547.1 |
| FAD/NAD(P)-binding protein [Streptomyces phyllanthi] | Streptomyces phyllanthi | 919 | 919 | 97% | 0.0 | 71.07% | 639 | WP_228031868.1 |
| FAD/NAD(P)-binding protein [Streptomyces sp. KM273126] | Streptomyces sp. KM273126 | 917 | 917 | 96% | 0.0 | 72.89% | 648 | WP_131571629.1 |
| hypothetical protein GCM10011579_018640 [Streptomyces albiflavescens] | Streptomyces albiflavescens | 917 | 917 | 98% | 0.0 | 73.55% | 711 | GGN56928.1 |
| FAD/NAD(P)-binding protein [Streptomyces albiflavescens] | Streptomyces albiflavescens | 916 | 916 | 98% | 0.0 | 73.55% | 648 | WP_229702725.1 |
| hypothetical protein ALECFALPRED_002312 [Alectoria fallacina] | Alectoria fallacina | 916 | 916 | 93% | 0.0 | 73.49% | 644 | CAF9923151.1 |
| FAD/NAD(P)-binding protein [unclassified Streptomyces] | unclassified Streptomyces | 915 | 915 | 98% | 0.0 | 71.16% | 652 | WP_330304262.1 |
| FAD/NAD(P)-binding protein [Streptomyces sp. NBC_01231] | Streptomyces sp. NBC_01231 | 915 | 915 | 98% | 0.0 | 71.03% | 644 | WSQ08710.1 |
| FAD/NAD(P)-binding protein [Streptomyces sp. NBC_00038] | Streptomyces sp. NBC_00038 | 914 | 914 | 98% | 0.0 | 70.55% | 654 | WP_266826772.1 |
| FAD/NAD(P)-binding protein [Streptomyces] | Streptomyces | 913 | 913 | 98% | 0.0 | 71.32% | 652 | WP_168491017.1 |
| uncharacterized protein N7509_001446 [Penicillium cosmopolitanum] | Penicillium cosmopolitanum | 909 | 909 | 96% | 0.0 | 71.63% | 639 | XP_056491878.1 |

**Figure S1.** blastp analysis of NpaA homologs. Using NpaA as a query gives a clear mix of both bacterial and fungal homolog hits with ~70% percent identity, which is remarkably high given they are members of different kingdoms of life.

| | | | | | | | | |
|---|---|---|---|---|---|---|---|---|
| uncharacterized protein N7509_001446 [Penicillium cosmopolitanum] | Penicillium ... | 909 | 909 | 96% | 0.0 | 71.63% | 639 | XP_056491878.1 |
| FAD/NAD(P)-binding protein [Streptomyces sp. DG2A-72] | Streptomyc... | 909 | 909 | 96% | 0.0 | 72.13% | 637 | WP_301979684.1 |
| uncharacterized protein N7481_010193 [Penicillium waksmanii] | Penicillium ... | 908 | 908 | 96% | 0.0 | 72.11% | 639 | XP_057120089.1 |
| FAD/NAD(P)-binding protein [Streptomyces albicerus] | Streptomyc... | 907 | 907 | 98% | 0.0 | 70.37% | 652 | WP_151480683.1 |
| FAD/NAD(P)-binding protein [Streptomyces sp. AS02] | Streptomyc... | 907 | 907 | 97% | 0.0 | 70.91% | 646 | WP_250093234.1 |
| FAD/NAD(P)-binding protein [Streptomyces sp. NBC_01136] | Streptomyc... | 907 | 907 | 98% | 0.0 | 71.67% | 652 | WST75471.1 |
| FAD/NAD(P)-binding protein [Streptomyces acidicola] | Streptomyc... | 907 | 907 | 97% | 0.0 | 71.50% | 640 | WP_152869882.1 |
| hypothetical protein NOR_03869 [Metarhizium rileyi RCEF 4871] | Metarhizium... | 907 | 907 | 92% | 0.0 | 72.30% | 667 | OAA44141.1 |
| FAD/NAD(P)-binding protein [Streptomyces sp. NBC_01236] | Streptomyc... | 906 | 906 | 98% | 0.0 | 69.56% | 664 | WSP70737.1 |
| FAD/NAD(P)-binding protein [Streptomyces sp. NBC_01235] | Streptomyc... | 906 | 906 | 96% | 0.0 | 72.06% | 637 | WSP86776.1 |
| hypothetical protein N0V93_002056 [Gnomoniopsis smithogilvyi] | Gnomoniop... | 905 | 905 | 93% | 0.0 | 70.49% | 666 | KAJ4397819.1 |
| FAD/NAD(P)-binding protein [Streptomyces sp. S3(2020)] | Streptomyc... | 904 | 904 | 96% | 0.0 | 70.84% | 638 | WP_171146718.1 |
| FAD-NAD(P)-binding protein [Metarhizium robertsii] | Metarhizium... | 904 | 904 | 91% | 0.0 | 73.17% | 660 | EXU97730.1 |
| FAD/NAD(P)-binding protein [Streptomyces cyaneochromogenes] | Streptomyc... | 904 | 904 | 97% | 0.0 | 70.60% | 646 | WP_126391900.1 |
| FAD/NAD(P)-binding protein [Streptomyces liliifuscus] | Streptomyc... | 904 | 904 | 96% | 0.0 | 71.04% | 689 | QQM47047.1 |
| FAD-NAD(P)-binding-domain-containing protein [Pestalotiopsis sp. NC0098] | Pestalotiops... | 903 | 903 | 94% | 0.0 | 69.64% | 642 | KAI0151355.1 |
| FAD/NAD(P)-binding protein [Streptomyces spinoverrucosus] | Streptomyc... | 903 | 903 | 97% | 0.0 | 70.00% | 638 | WP_196463046.1 |
| FAD/NAD(P)-binding protein [Streptomyces sp. CGMCC 4.7035] | Streptomyc... | 902 | 902 | 96% | 0.0 | 71.07% | 643 | WP_310781283.1 |
| hypothetical protein G6M90_00g113260 [Metarhizium brunneum] | Metarhizium... | 902 | 902 | 91% | 0.0 | 73.00% | 660 | QLI74210.1 |
| putative NAD(P)/FAD-binding protein YdhS [Streptomyces sp. V3I8] | Streptomyc... | 902 | 902 | 95% | 0.0 | 71.75% | 678 | MDQ1038927.1 |
| FAD/NAD(P)-binding protein [Streptomyces sp. ALI-76-A] | Streptomyc... | 902 | 902 | 96% | 0.0 | 70.20% | 648 | WP_286062116.1 |
| FAD(NAD)-dependent oxidoreductase [Metarhizium guizhouense ARSEF 977] | Metarhizium... | 902 | 902 | 91% | 0.0 | 73.42% | 660 | KID83344.1 |
| hypothetical protein KJ359_003818 [Pestalotiopsis sp. 9143b] | Pestalotiops... | 902 | 902 | 93% | 0.0 | 70.49% | 664 | KAI4598009.1 |
| FAD/NAD(P)-binding protein [Streptomyces fulvoviolaceus] | Streptomyc... | 902 | 902 | 98% | 0.0 | 70.87% | 645 | WP_261726153.1 |
| hypothetical protein MHUMG1_05852 [Metarhizium humberi] | Metarhizium... | 901 | 901 | 91% | 0.0 | 73.17% | 660 | KAH0596732.1 |
| hypothetical protein E5D57_012337 [Metarhizium anisopliae] | Metarhizium... | 901 | 901 | 91% | 0.0 | 72.95% | 659 | KAF5121867.1 |
| putative NAD(P)/FAD-binding protein YdhS [Streptomyces umbrinus] | Streptomyc... | 901 | 901 | 97% | 0.0 | 70.58% | 669 | MDQ1029621.1 |
| hypothetical protein GCM10010383_07870 [Streptomyces lomondensis] | Streptomyc... | 900 | 900 | 96% | 0.0 | 70.44% | 828 | GGW82146.1 |
| FAD/NAD(P)-binding protein [Streptomyces] | Streptomyces | 900 | 900 | 98% | 0.0 | 69.56% | 668 | WP_099504500.1 |
| hypothetical protein IMSHALPRED_001622 [Imshaugia aleurites] | Imshaugia a... | 900 | 900 | 93% | 0.0 | 70.73% | 633 | CAF9939763.1 |
| FAD/NAD(P)-binding protein [Streptomyces phyllanthi] | Streptomyc... | 899 | 899 | 94% | 0.0 | 71.25% | 621 | MPY45702.1 |
| FAD/NAD(P)-binding protein [Streptomyces sp. NBC_00299] | Streptomyc... | 899 | 899 | 97% | 0.0 | 70.38% | 639 | WP_328881978.1 |
| FAD/NAD(P)-binding protein [Streptomyces umbrinus] | Streptomyc... | 899 | 899 | 96% | 0.0 | 71.20% | 632 | WP_307531691.1 |
| FAD/NAD(P)-binding protein [Streptomyces bluensis] | Streptomyc... | 899 | 899 | 96% | 0.0 | 71.61% | 648 | WP_189872334.1 |
| putative NAD(P)/FAD-binding protein YdhS [Streptomyces phaeochromogenes] | Streptomyc... | 899 | 899 | 96% | 0.0 | 70.35% | 716 | MDQ0952824.1 |
| FAD/NAD(P)-binding protein [Streptomyces sp. NBC_00286] | Streptomyc... | 899 | 899 | 98% | 0.0 | 69.81% | 649 | WP_328771982.1 |
| FAD/NAD(P)-binding protein [Streptomyces sp. NBC_01352] | Streptomyc... | 899 | 899 | 96% | 0.0 | 70.96% | 644 | WP_329341151.1 |
| FAD/NAD(P)-binding protein [Streptomyces sp. V4I2] | Streptomyc... | 898 | 898 | 98% | 0.0 | 69.80% | 644 | WP_307674499.1 |
| FAD/NAD(P)-binding protein [Streptomyces fulvoviolaceus] | Streptomyc... | 898 | 898 | 98% | 0.0 | 70.72% | 645 | WP_078655513.1 |
| FAD/NAD(P)-binding protein [Streptomyces phaeochromogenes] | Streptomyc... | 898 | 898 | 96% | 0.0 | 70.89% | 630 | WST00206.1 |
| uncharacterized protein MAM_06969 [Metarhizium album ARSEF 1941] | Metarhizium... | 897 | 897 | 91% | 0.0 | 72.07% | 663 | XP_040676324.1 |
| FAD(NAD)-dependent oxidoreductase [Metarhizium majus ARSEF 297] | Metarhizium... | 896 | 896 | 91% | 0.0 | 73.04% | 659 | KID96183.1 |
| FAD/NAD(P)-binding protein [Streptomyces sp. NBC_00287] | Streptomyc... | 896 | 896 | 94% | 0.0 | 72.12% | 640 | WP_328871052.1 |
| putative NAD(P)/FAD-binding protein YdhS [Streptomyces umbrinus] | Streptomyc... | 895 | 895 | 96% | 0.0 | 70.20% | 669 | MCR3729812.1 |

**Figure S1. (continued)** blastp analysis of NpaA homologs. Using NpaA as a query gives a clear mix of both bacterial and fungal homolog hits with ~70% percent identity, which is remarkably high given they are members of different kingdoms of life.

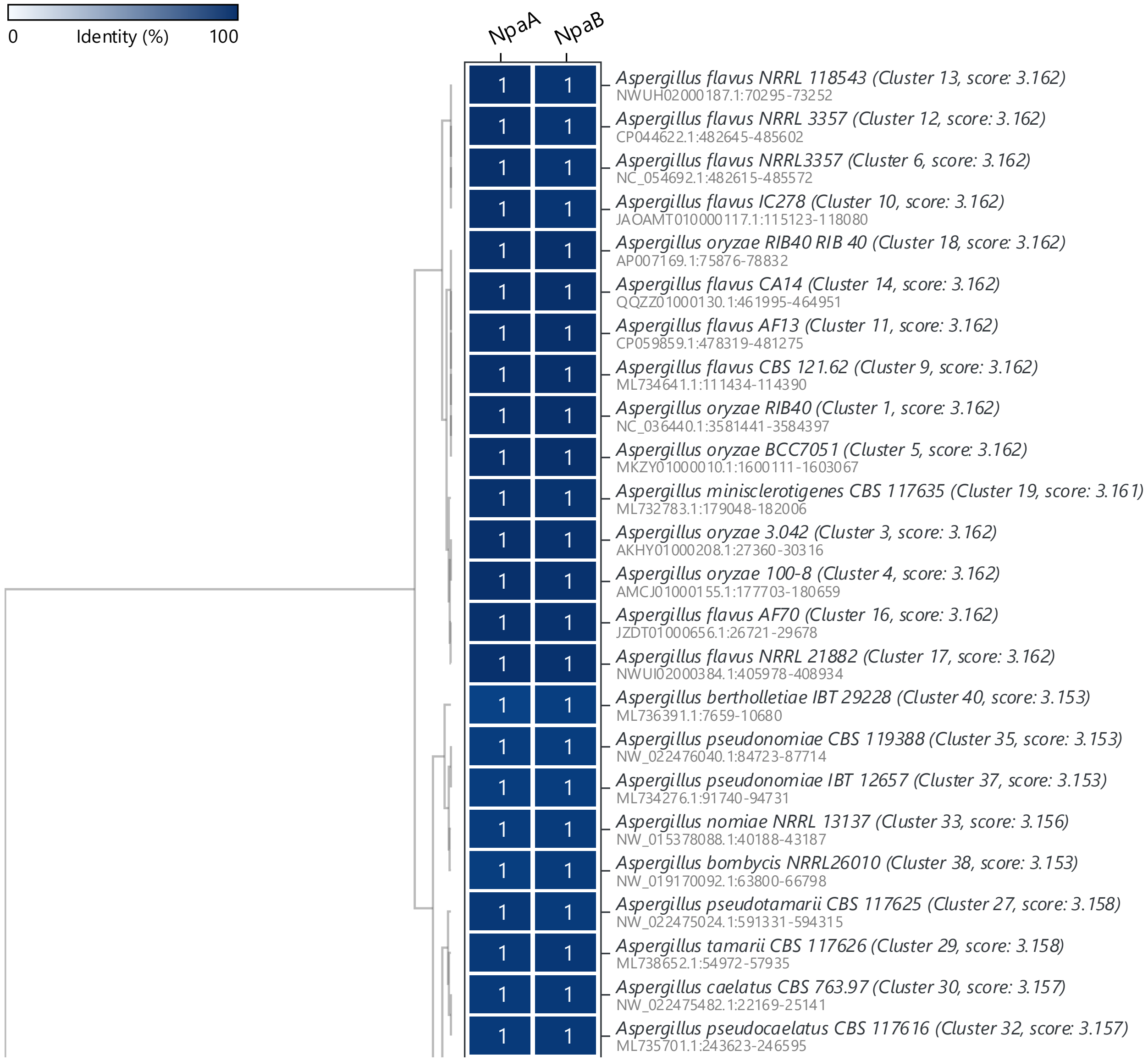


**Figure S2**. **(Continued)** Distribution of *npa* biosynthetic gene cluster in fungi from the NCBI database identified by cblaster.[7]

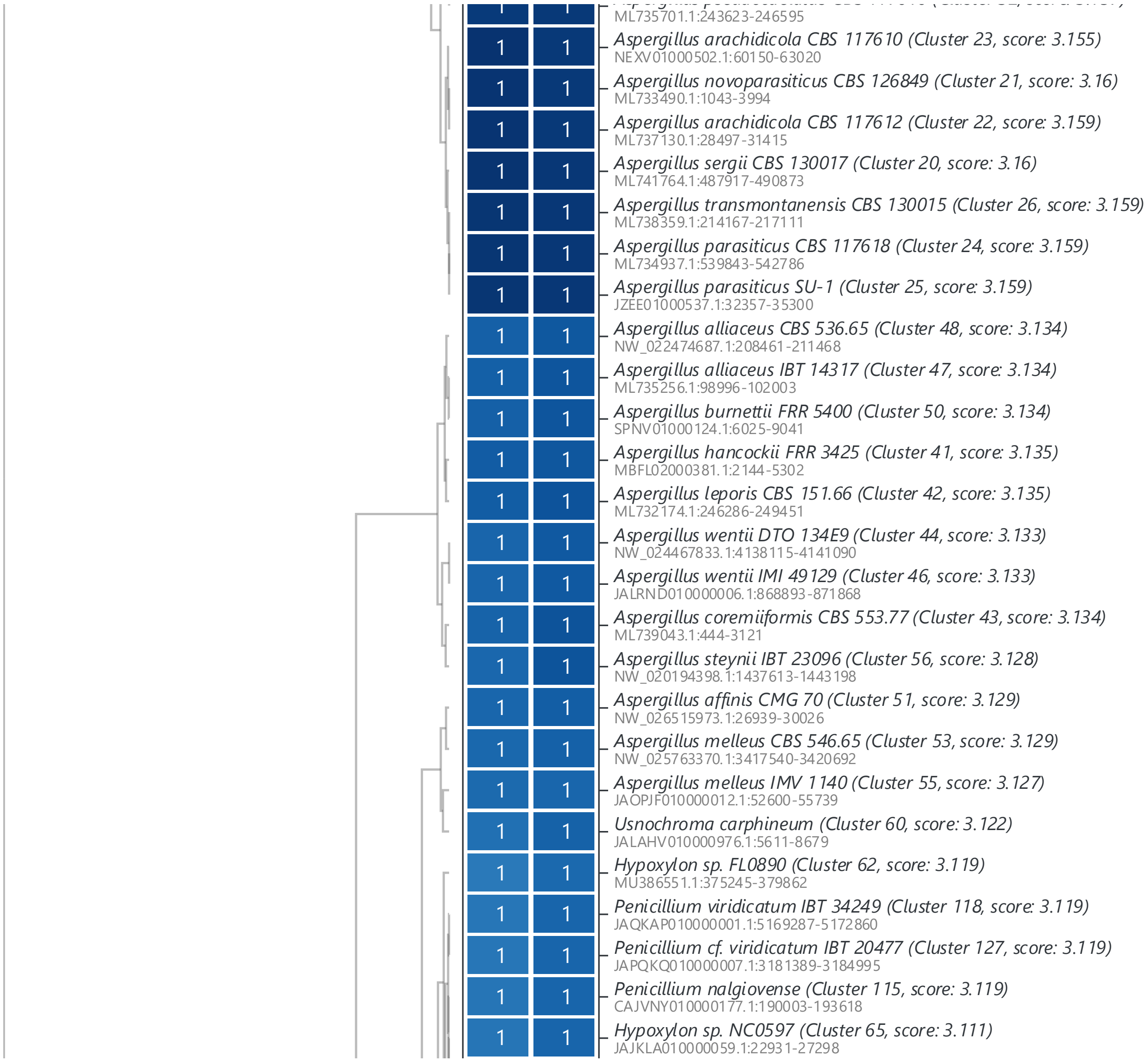


**Figure S2. (Continued)** Distribution of *npa* biosynthetic gene cluster in fungi from the NCBI database identified by cblaster.[7]

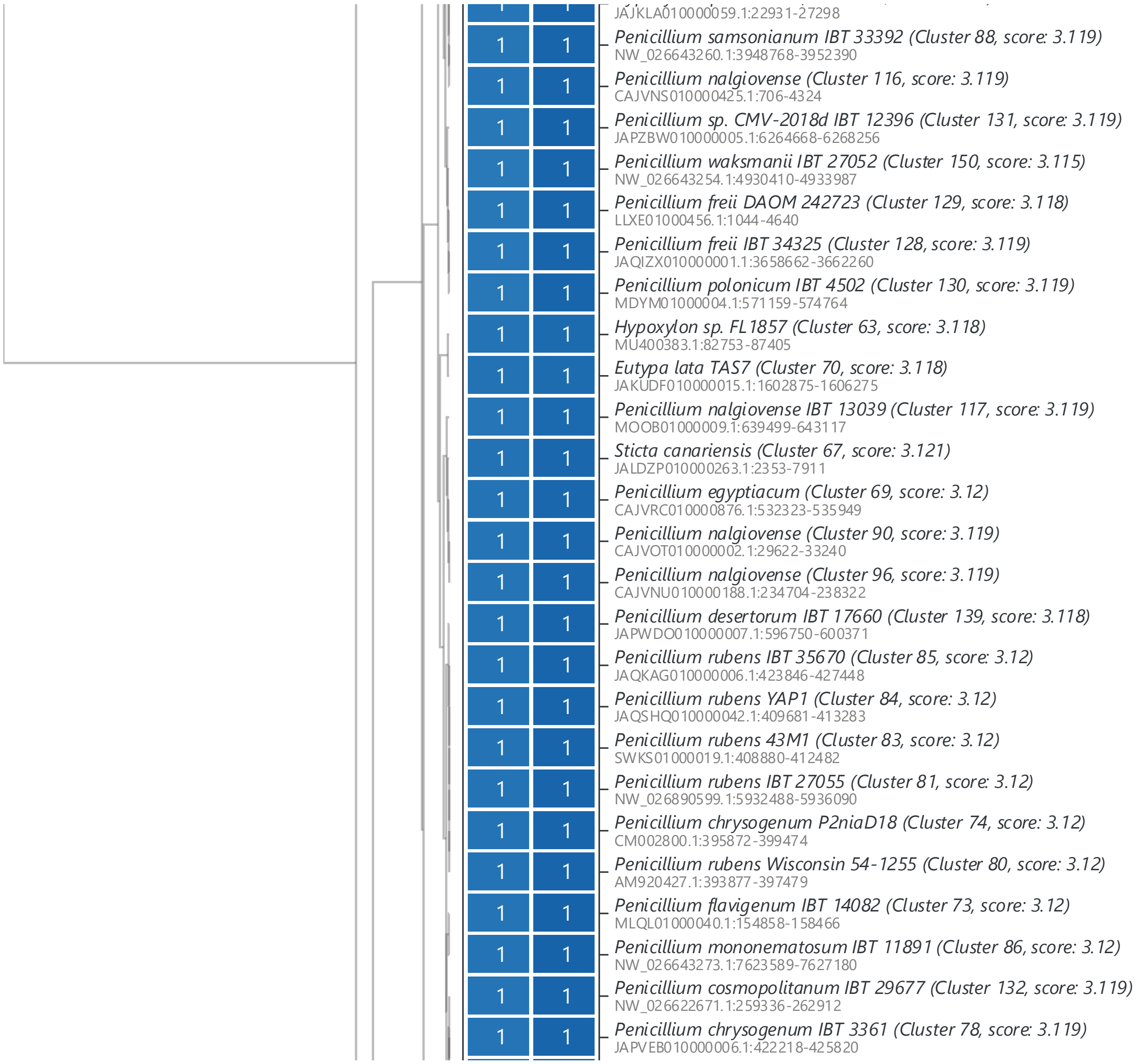


**Figure S2**. **(Continued)** Distribution of *npa* biosynthetic gene cluster in fungi from the NCBI database identified by cblaster.[7]

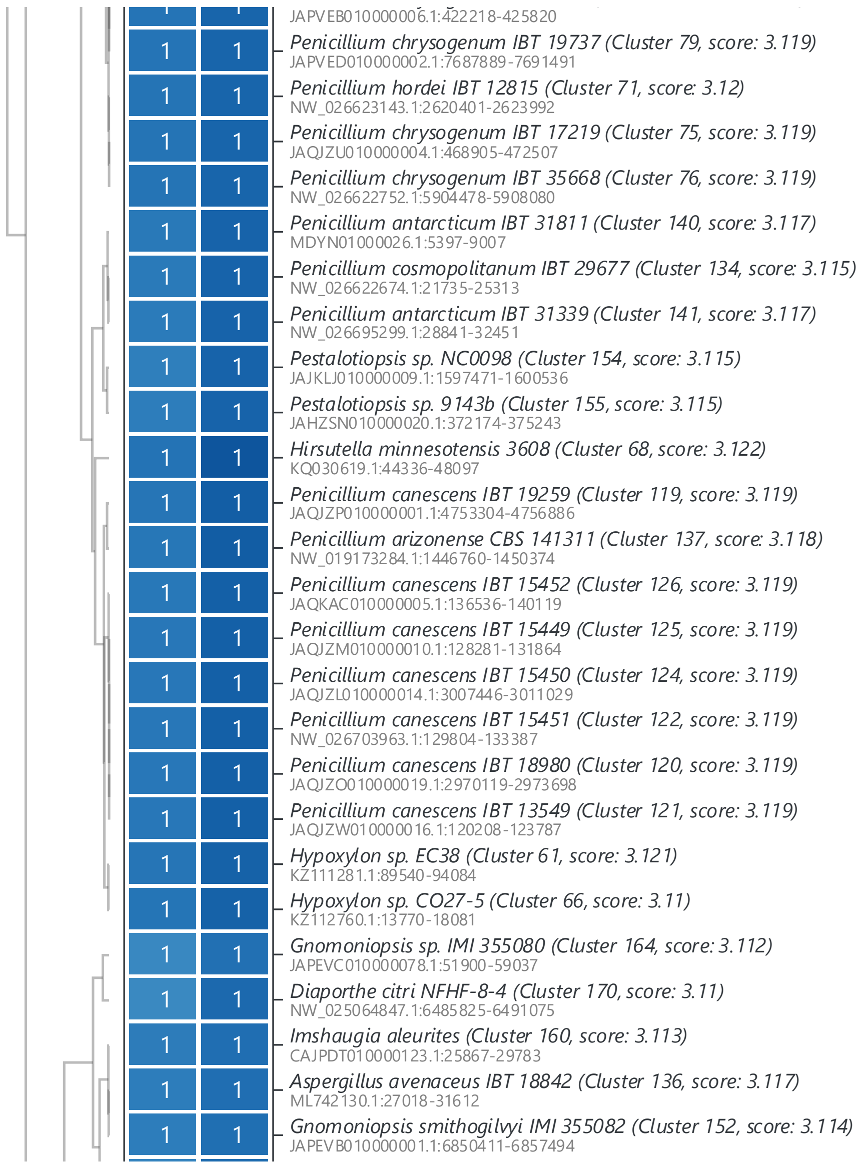


**Figure S2**. **(Continued)** Distribution of *npa* biosynthetic gene cluster in fungi from the NCBI database identified by cblaster.[7]

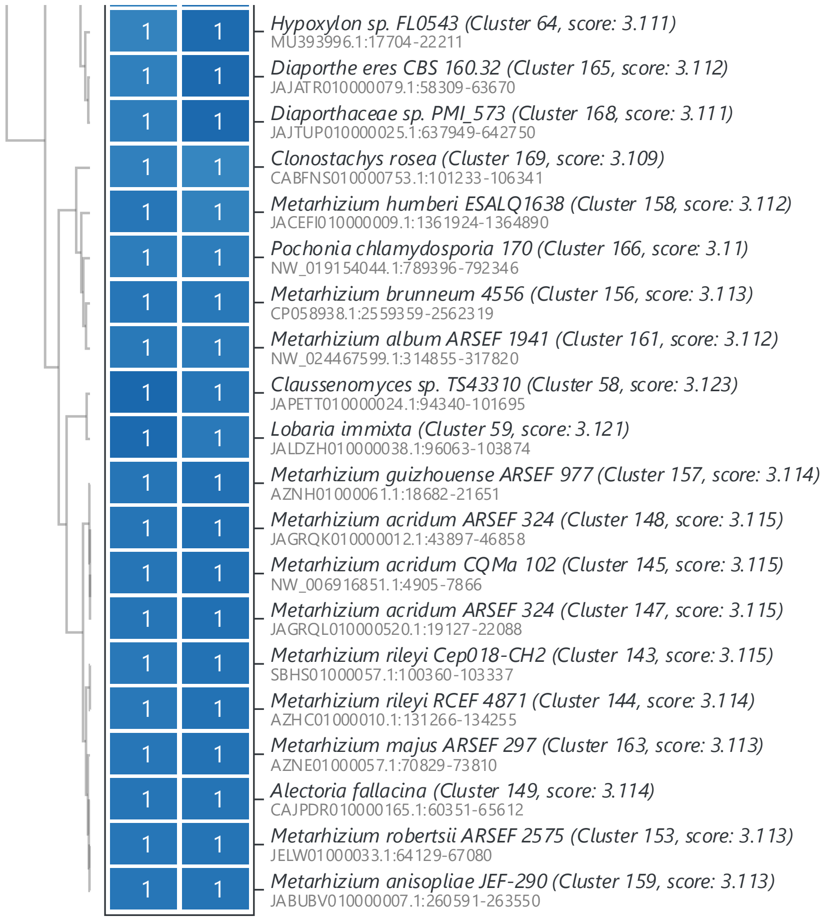


**Figure S2**. Distribution of *npa* biosynthetic gene cluster in fungi from the NCBI database identified by cblaster.[7]

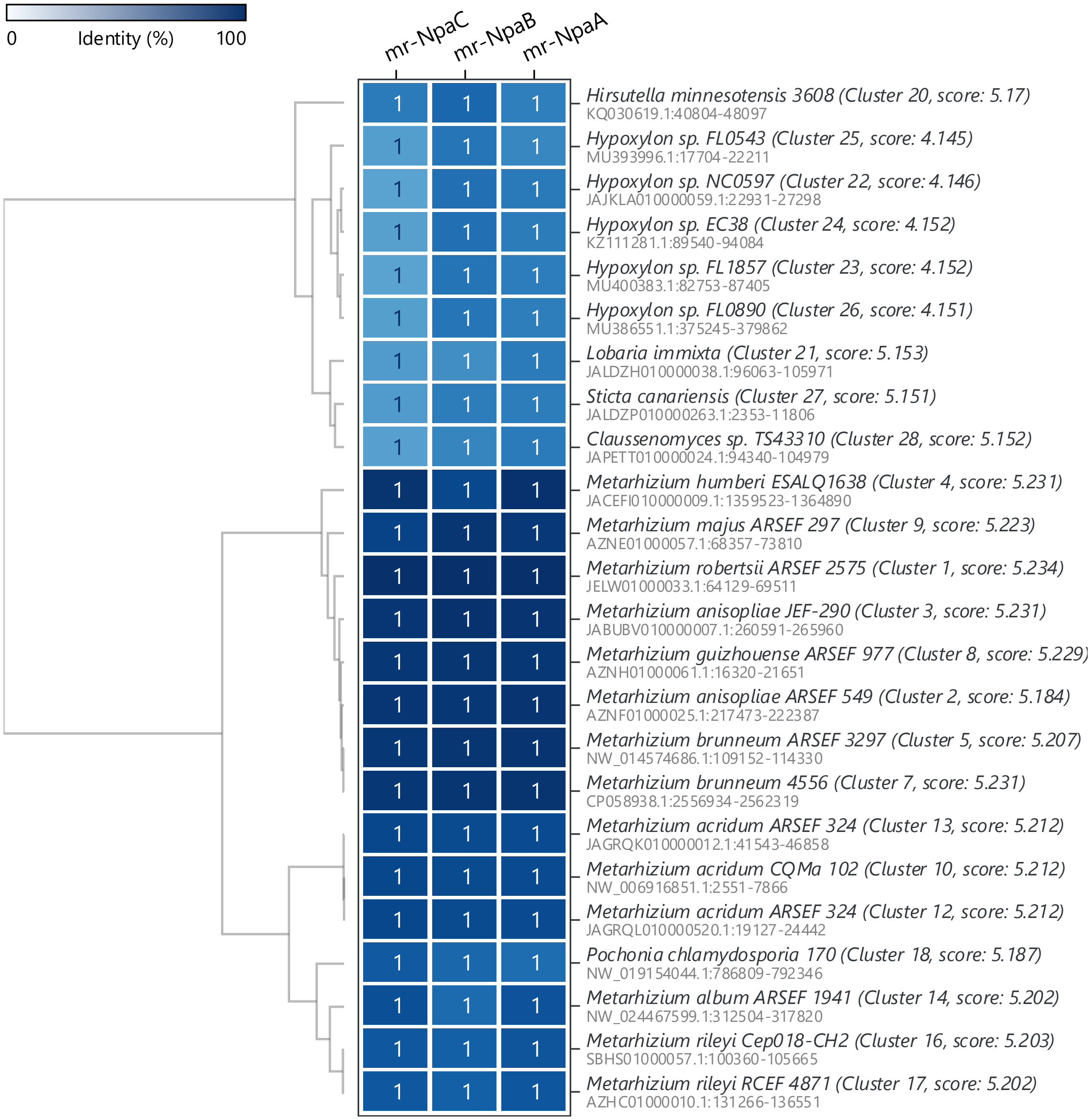


**Figure S3**. Example *npa* clusters encoding a third conserved gene *npaC* identified by cblaster.[7]

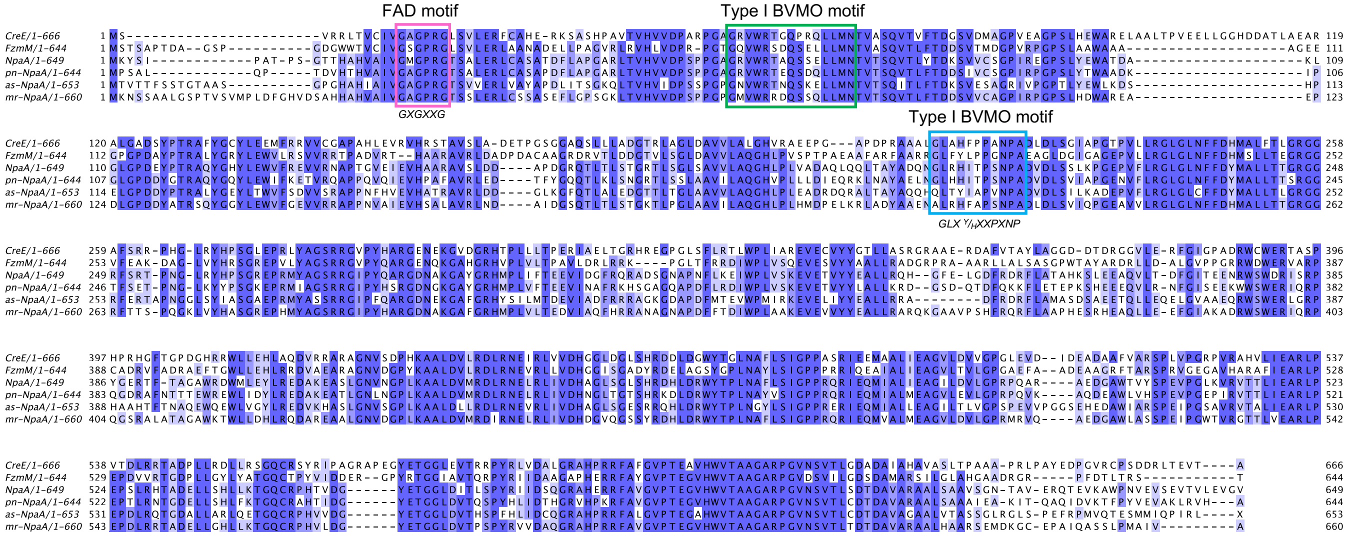


**Figure S4.** Multiple sequence alignments of CreE homologs including NpaA and FmzM. The T-Coffee server[8] was used for sequence alignment and the software Jalview[9]) was used for visualization. Residues are colored blue depending on percent identity. The residues surrounded by the pink rectangle depict the FAD binding motif, green rectangle is the type I BVMO motif involved in FAD binding, blue rectangle is the type I BVMO motif involved in NADPH binding. As similar to CreE, NpaA homologs all have type I BVMO motif.[10]

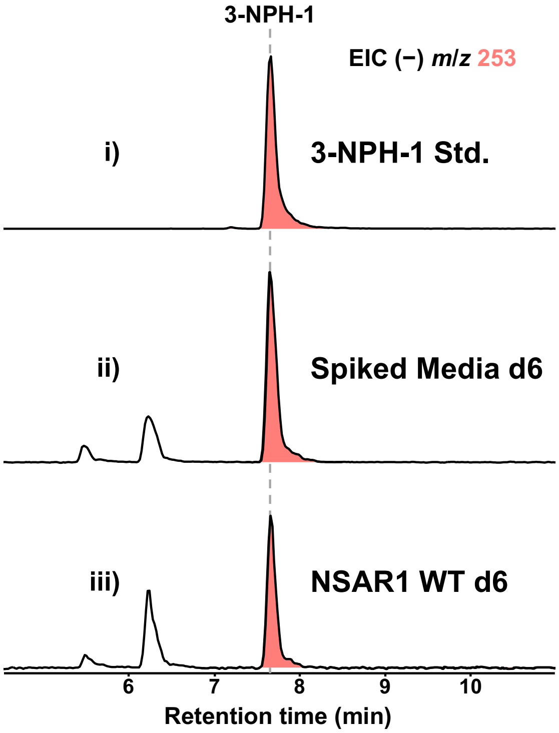


**Figure S5.** Validation of 3-NPH derivatization strategy for *in vivo* detection of **1** via **3-NPH-1**. (i) LC-MS analysis of authentic **3-NPH-1** standard. (ii). To ensure **1** could be efficiently extracted and detected from aqueous culture media, Nakamura media (200 mL) was spiked with 5 mg of **1** standard and subjected to identical culture conditions used to culture *A. oryzae* NSAR1 (30 ˚C shaking at 220 rpm). After 6 days, the culture was harvested as described above and **1** was able to be recovered and detected in (ii) as **3-NPH-1**. (iii) Detection of **1** in *A. oryzae* NSAR1 WT strain after 6 days using the culture conditions and detection protocol described above.

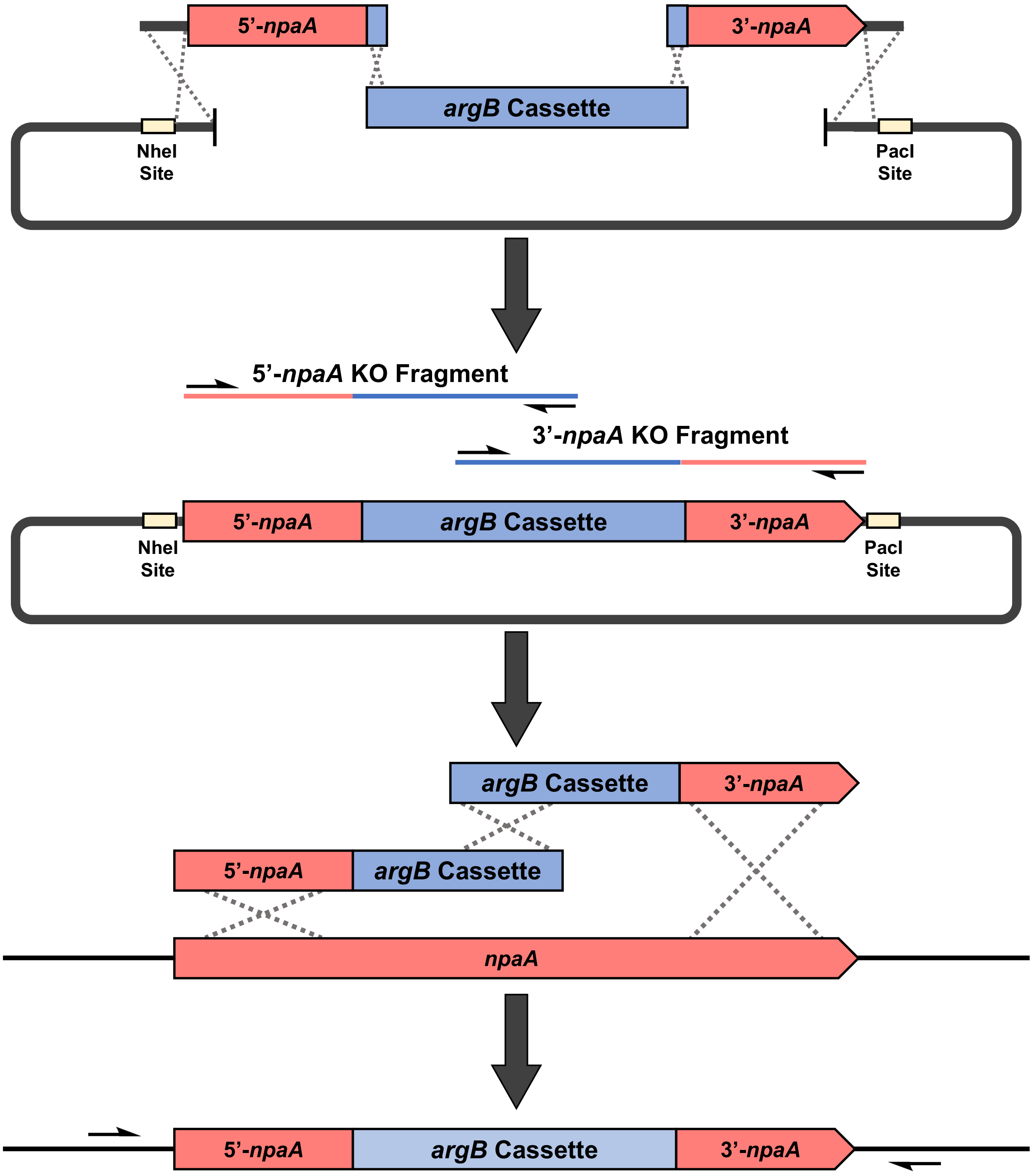


**Figure S6.** Scheme of *npaA* disruption cassette design for *npaA* KO in *A. oryzae* NSAR1.

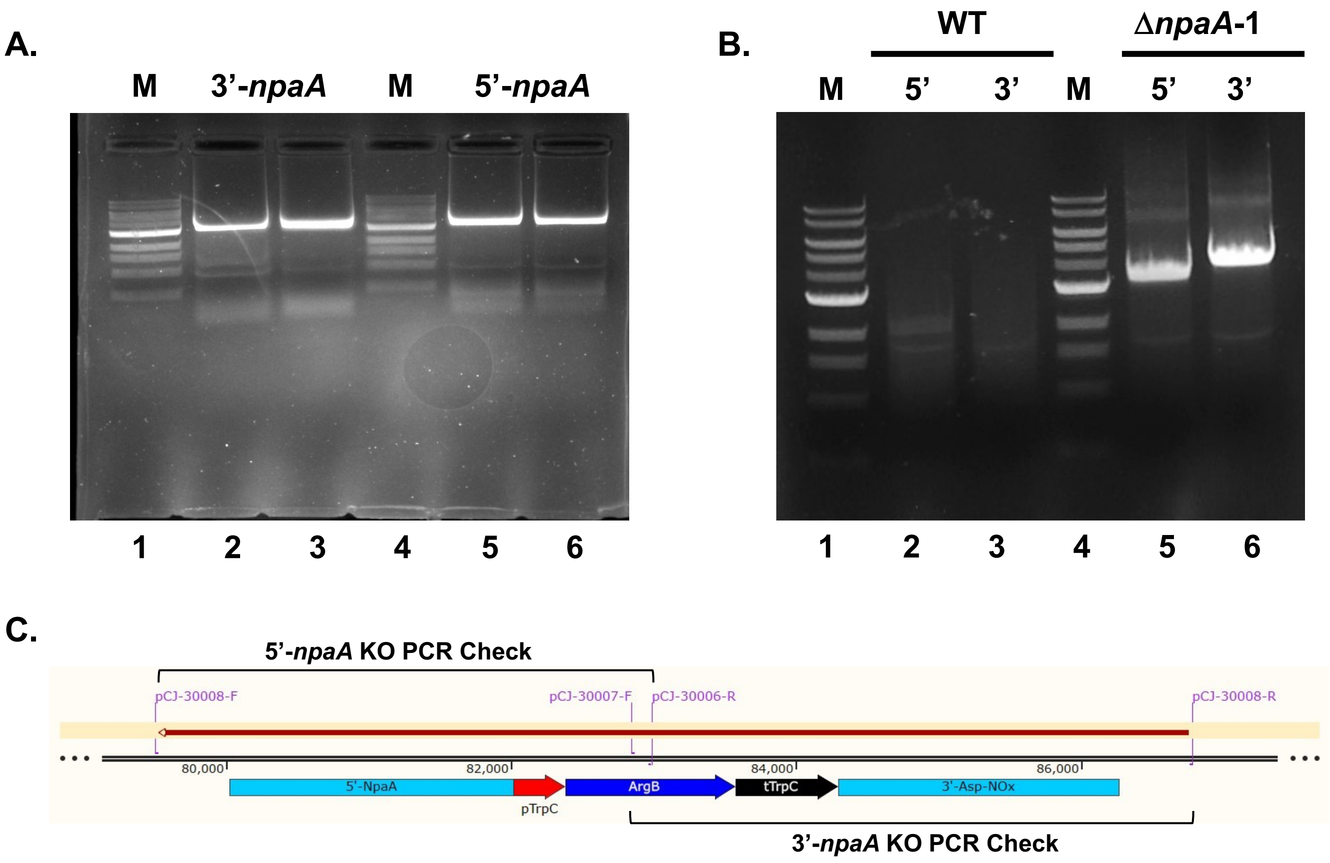


**Figure S7.** Amplification of *npaA* bipartite knockout fragments. **A.** PCR amplification of *npaA* 3' and 5' fragments for split-marker knockout approach of *npaA* in *A. oryzae* NSAR1. Lane 1, 1kb DNA ladder; lanes 2-3, 3'-*npaA* KO fragment; lane 4, 1kb DNA ladder; lanes 5-6, 5'-*npaA* KO fragment. **B.** Diagnostic PCR check for integration of ArgB disruption cassette. Amplification is only possible with successful integration. Lane 1, 1kb DNA ladder; lane 2, 5'-*npaA* KO check (amplified from pCJ-30008-F / pCJ-30006-R); lane 3, 3'-*npaA* KO check (amplified from pCJ-30007-F / pCJ-30008-R); lane 4, 1kb DNA ladder; lane 5, 5'-*npaA* KO check (amplified from pCJ-30008-F / pCJ-30006-R); and lane 6, 3'-*npaA* KO check (amplified from pCJ-30007-F / pCJ-30008-R). **C.** Scheme of fragments amplified when checking for *npaA* disruption used in **B**.

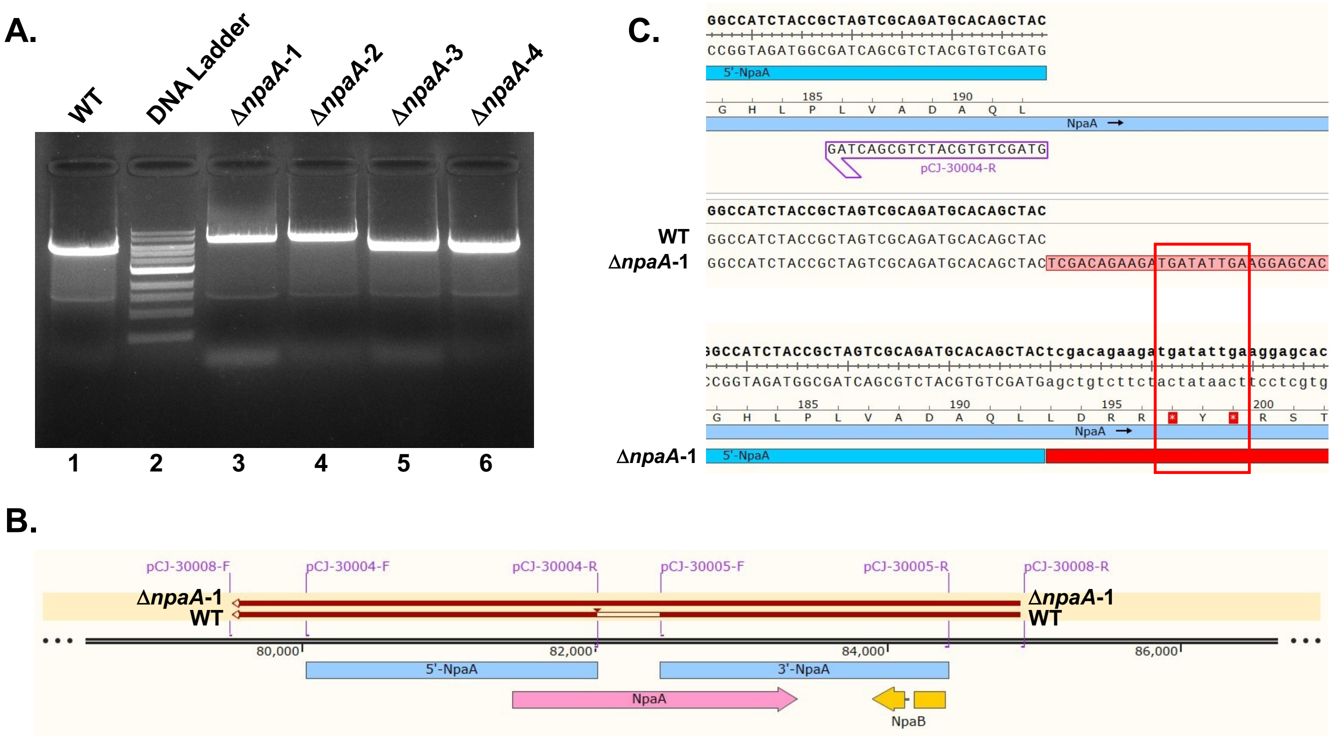


**Figure S8.** PCR and sequencing verification of successful *npaA* KO in *A. oryzae* NSAR1. **A.** PCR products from colonies of four knockout attempts (lanes 3-6) versus WT (lane 1). Increased size of amplicons in Δ*npaA*-1 and Δ*npaA*-2 (lanes 3 and 4) demonstrates integration compared to WT or unsuccessful integration in colonies Δ*npaA*-3 and Δ*npaA*-4 (lanes 5 and 6). **B.** Overview of sequencing data indicating integration of disruption cassette in Δ*npaA*-1. **C.** Detailed view of Δ*npaA*-1 sequencing results. The *npaA* gene was successfully disrupted evidenced by the insertion of stop codons (red box).

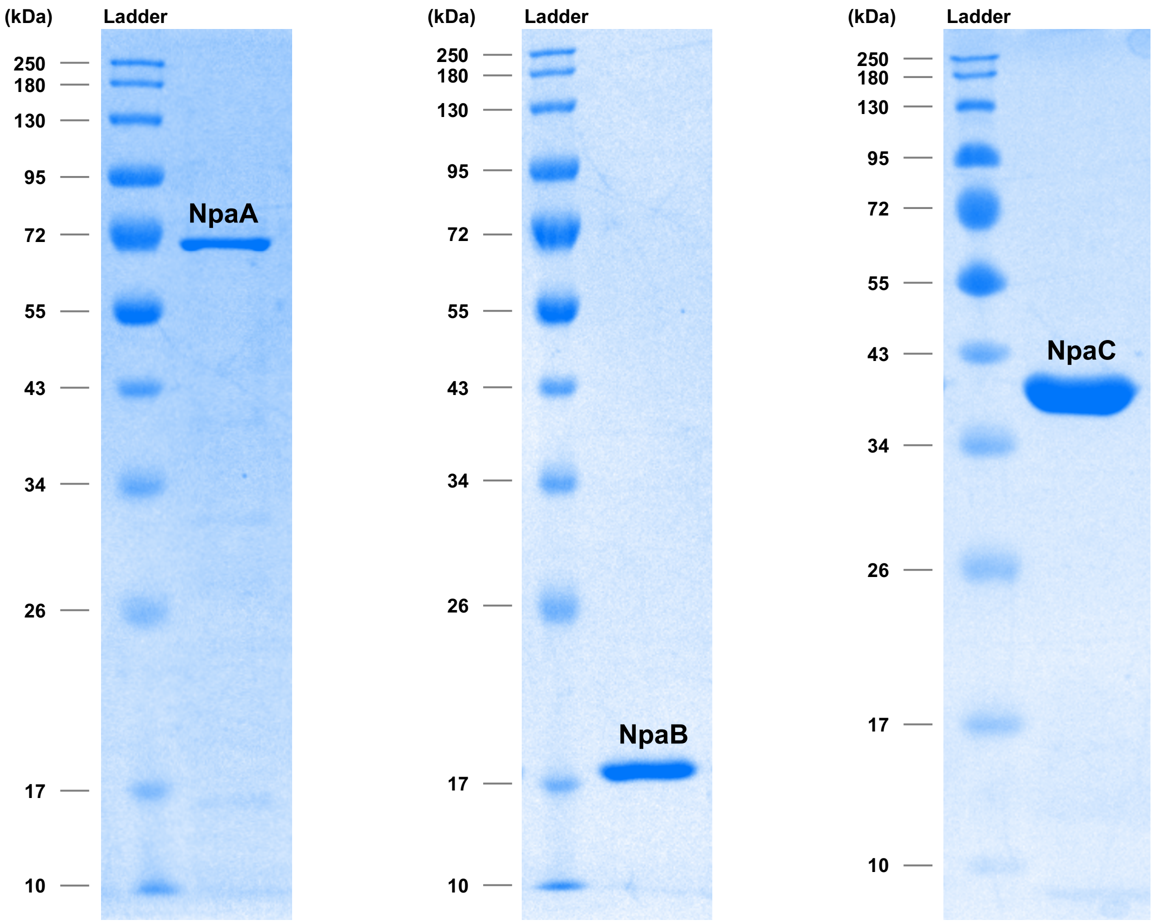


**Figure S9**. SDS−PAGE analyses of NpaA, NpaB, and mr-NpaC expressed and purified from *E. coli*.

**A.**

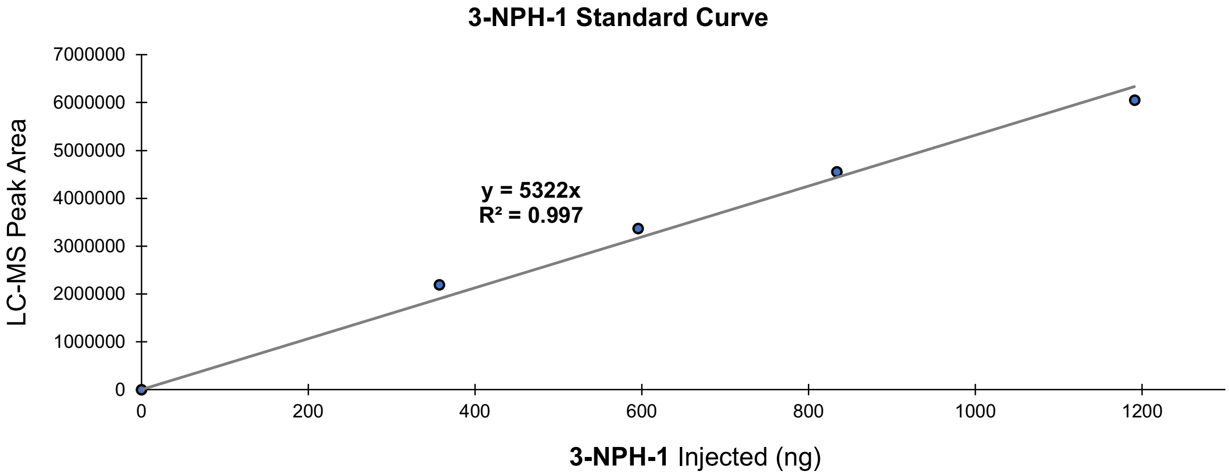


**B.**

| WT NSAR1 d6 3-NPH-1 Area | 3-NPH-1 (ng) | Dilution Factor (DF) | DF Corr. | 200mL Cx Extracted | ng/L | 3-NPH-1 mg/L |
|---|---|---|---|---|---|---|
| 405363 | 76.17 | 10 | 761.67 | 5 | 3808.37 | 3.81 |

**Figure S10**. Standard curve of **3-NPH-1** to estimate the production of **1** in *A. oryzae* NSAR1. **A.** Standard curve generated from 2 μL, 3 μL, 5 μL, 7 μL, and 10 μL injections of derivatized 1 mM **1** standard. **B.** Dilution-corrected approximate quantification (mg/L) of **3-NPH-1** calculated from standard curve in **A.**

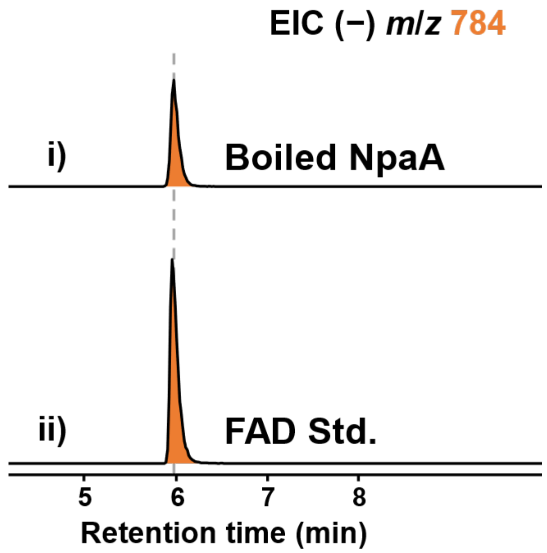


**Figure S11.** Identification of FAD as cofactor of NpaA. LC-MS analysis of boiled NpaA supernatant (i) compared with authentic FAD standard (ii).

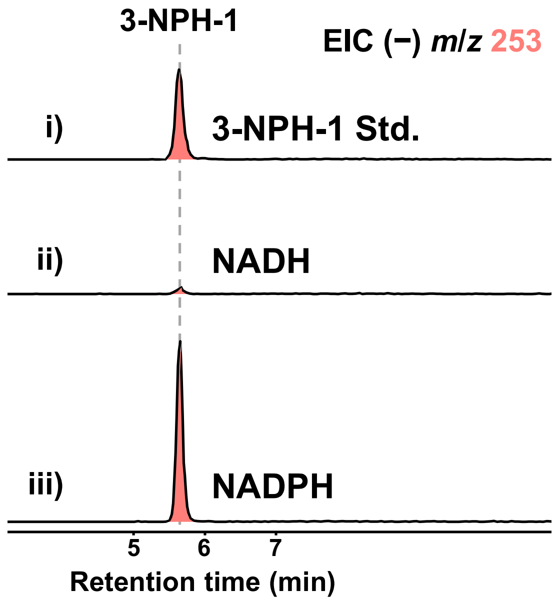


**Figure S12.** Identification of NADPH as native co-substrate of NpaA. LC-MS analysis 3-NPA production monitored by **3-NPH-1** of *in vitro* reaction of NpaA with NADH (ii) vs. NADPH (iii) in comparison to authentic **3-NPH-1** standard (i).

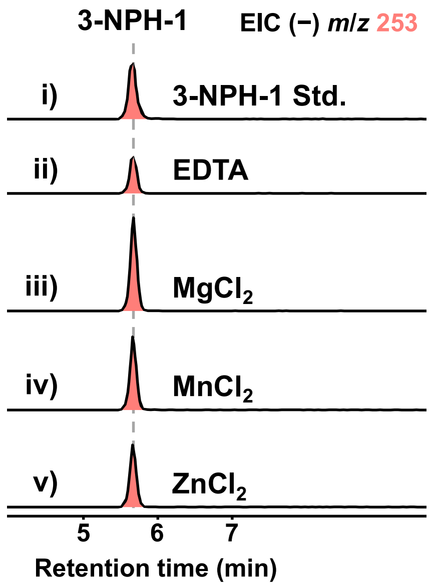


**Figure S13.** Screening EDTA-dialyzed NpaB with divalent metals. Comparison of 3-NPH-1 production by NpaA and NpaB (dialyzed with 1mM EDTA) after 15 minutes with (iii) $MgCl_2$, (iv) $MnCl_2$, or (v) $ZnCl_2$ monitored by LC-MS. Results demonstrate that all divalent metals tested accelerate the decarboxylation in contrast to EDTA-treated control (ii), with $MgCl_2$ showing the greatest effect.

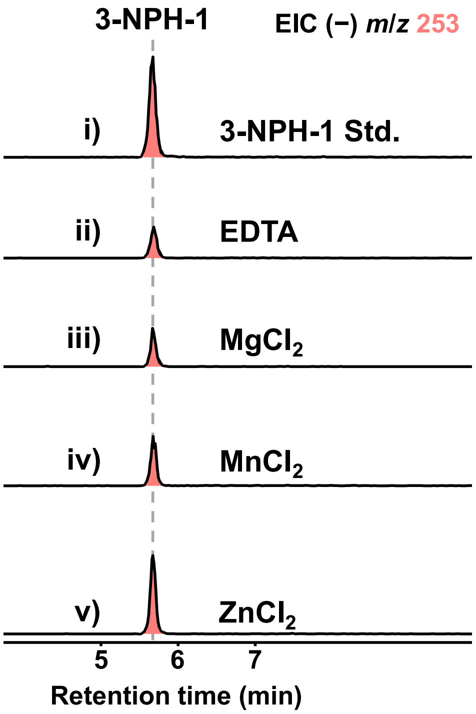


**Figure S14.** Non-enzymatic decarboxylation in the presence of boiled NpaB and screened divalent metals. Boiled EDTA-dialyzed NpaB was screened with the same divalent metals in **Fig. S12**. Non-enzymatic decarboxylation was increased most in the presence of $ZnCl_2$, (v).

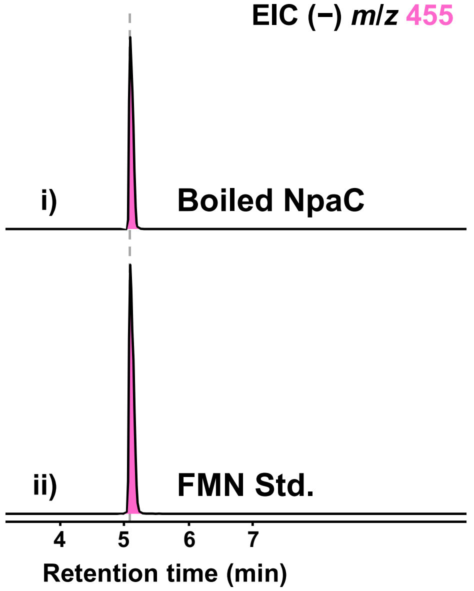


**Figure S15.** Identification of FMN as cofactor of mr-NpaC. LC-MS analysis of boiled mr-NpaC supernatant (i) compared with authentic FMN standard (ii).

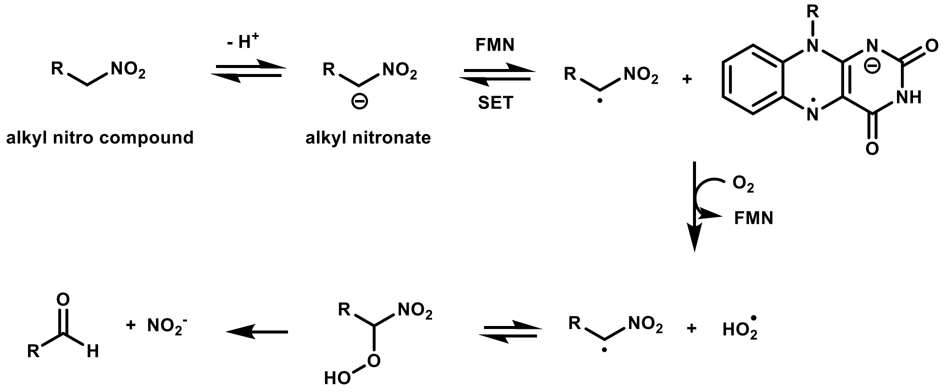


**Figure S16.** Representative example of NMO oxidation yielding carbonyl compounds and nitrite. General reactivity of nitronate monooxygenase. Single electron transfer between alkyl nitronate and FMN oxidizes alkyl nitronate and reduces FMN. Then molecular oxygen oxidizes FMN and resulting superoxide and alkyl radical undergo radical recombination yielding the alpha-peroxynitro alkane. This species decays resulting in carbonyl formation and release of nitrite.[11]

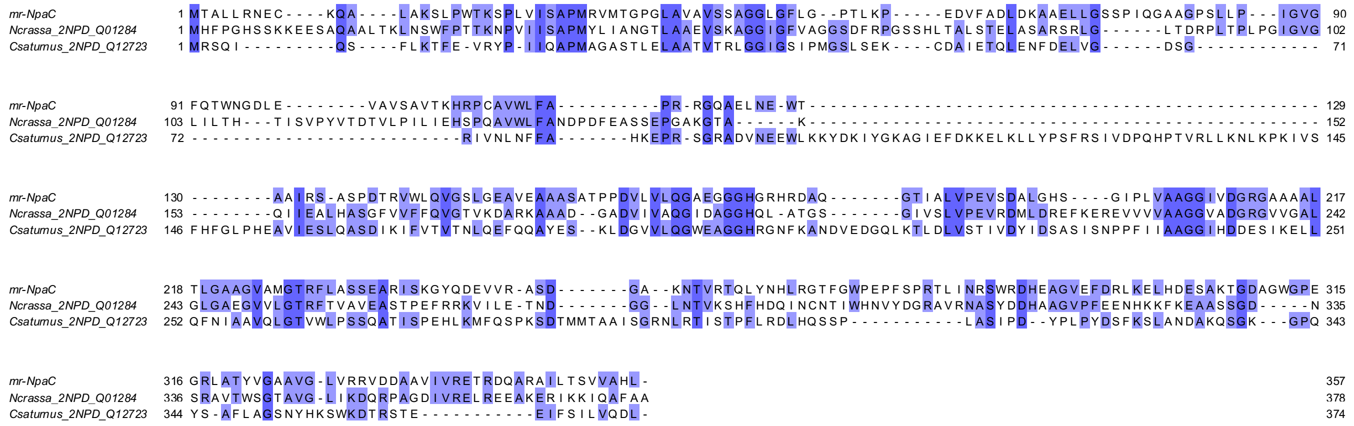


**Figure S17.** Sequence alignment of mr-NpaC with characterized fungal NMOs. The T-Coffee server[8] was used for sequence alignment and the software Jalview[9] was used for visualization. Residues are colored blue depending on percent identity. Two classes of NMOs (Class I and Class II) were established based on their distinct differences inferred from bioinformatic, mechanistic, and structural studies.[12] In fungi, NMOs from both Class I and Class II have been characterized. Recently, a Class I NMO from the yeast *Cyberlindnera saturnus* (cs-NMO) was characterized, while the only characterized member of Class II NMOs is from the filamentous fungi, *Neurospora crassa* (mc-NMO). Comparing mr-NpaC to characterized members of each class in the alignment above suggests mr-NpaC is a member of Class II NMOs.

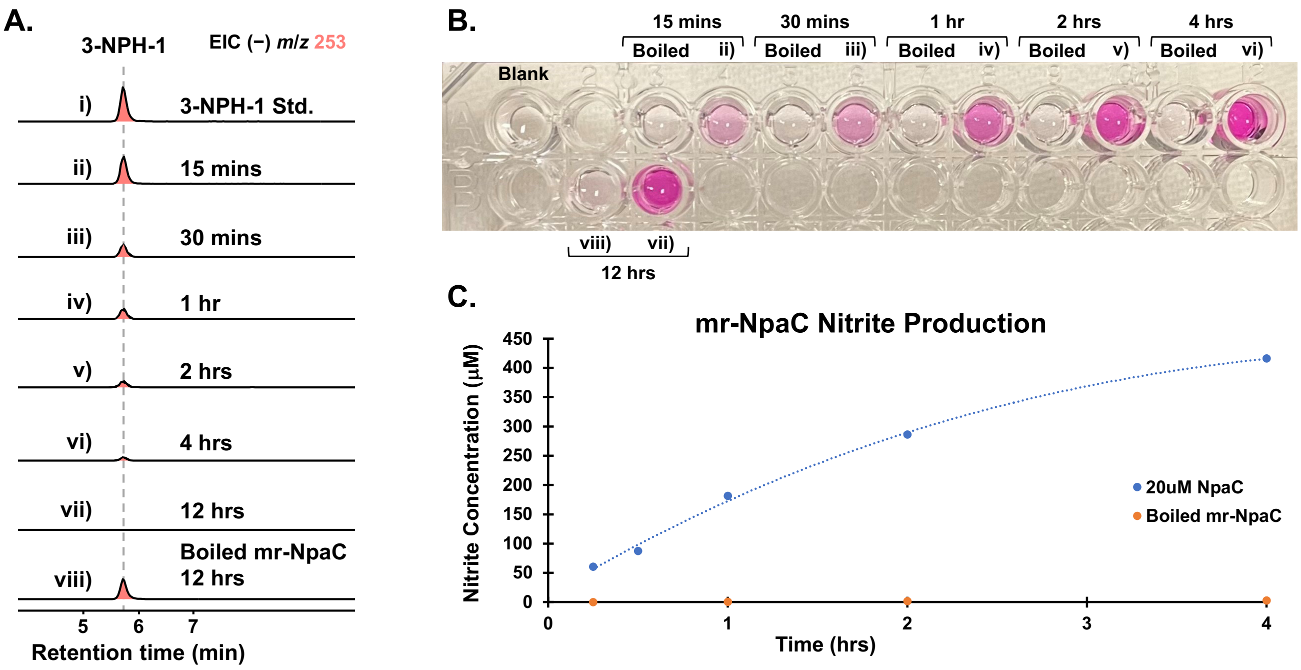


**Figure S18.** Time-dependent experiment of 3-NPA consumption and nitrite production by mr-NpaC. **A.** LC-MS monitoring of **3-NPH-1** (derivatized 3-NPA) at 15 mins (ii), 30 minutes (iii), 1 hr (iv), 2 hrs (v), 4 hrs (vi), and 12 hrs (vii) vs. boiled mr-NpaC control (viii) and **3-NPH-1** authentic standard (i). **B.** Detection of nitrite production via Griess test corresponding to *in vitro* timepoints shown in **A. C.** Quantification of nitrite production calculated from Griess test shown in **B**.

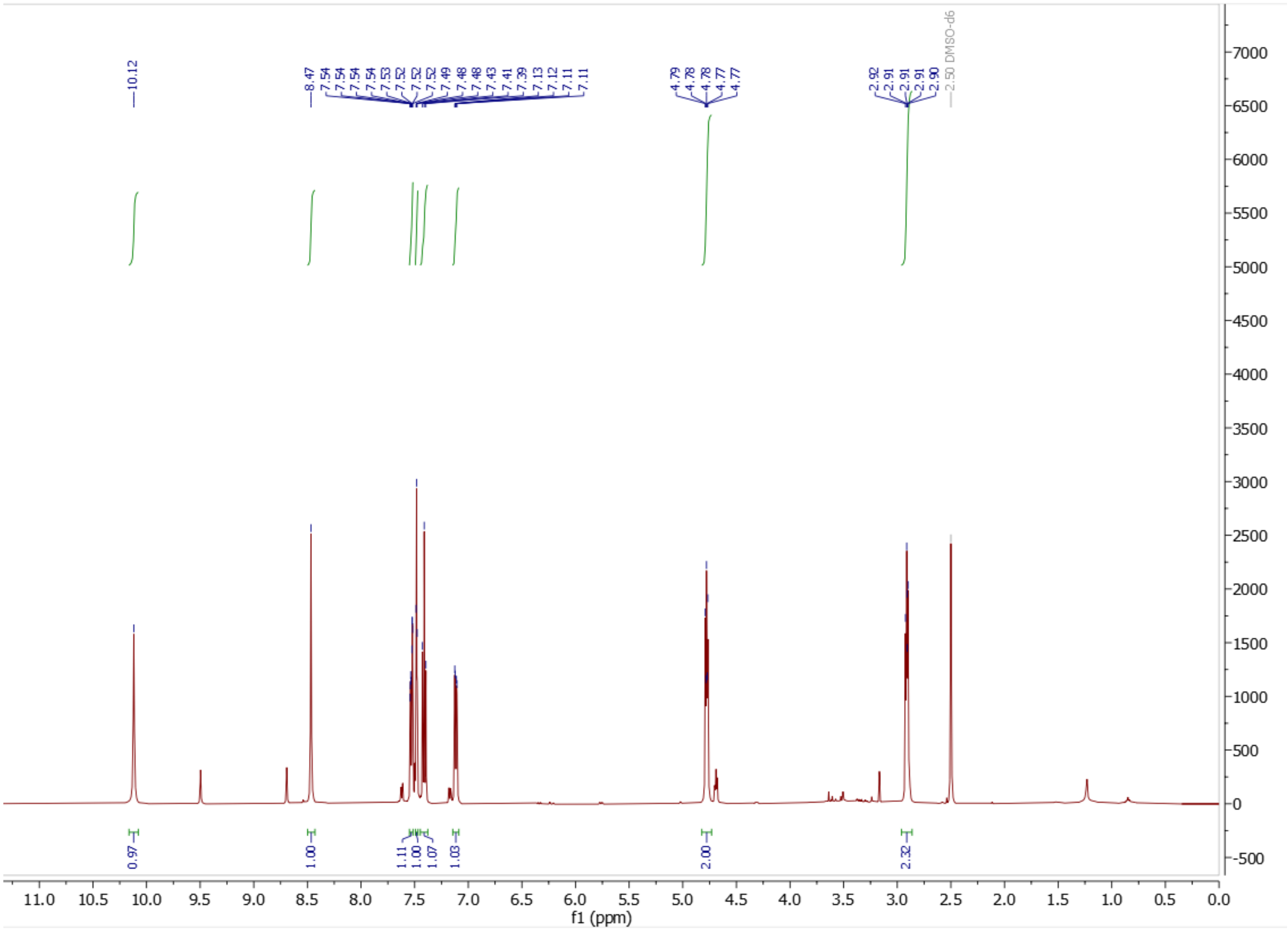


**Figure S19.** $^{1}H$ NMR spectrum of compound **3-NPH-1** in DMSO (500 MHz).

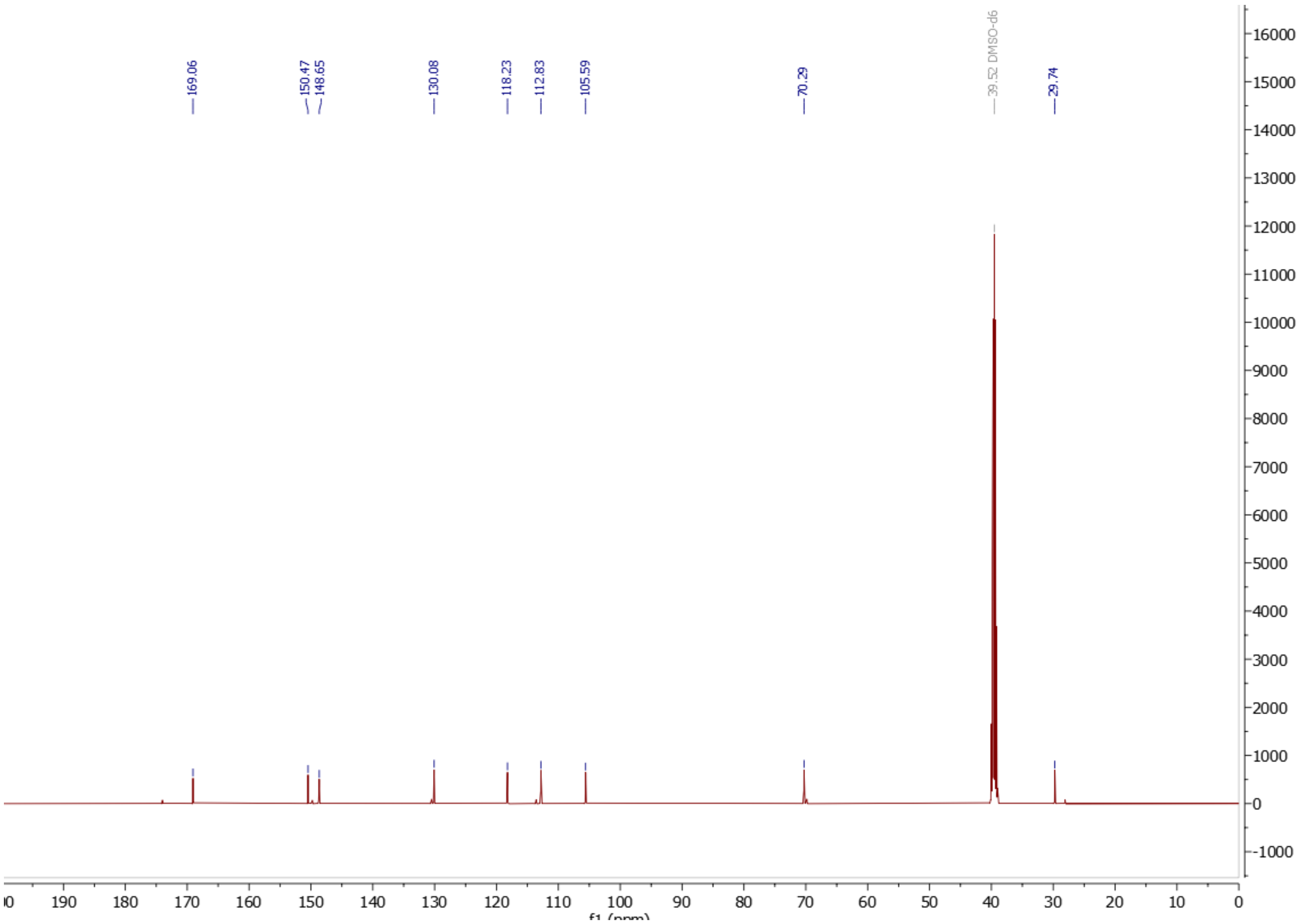


**Figure S20.** $^{13}$C NMR spectrum of compound **3-NPH-1** in DMSO (125 MHz).

#### Sequence Information

**DNA sequence of *npaA* homolog *as-npaA* gene.**

*as-npaA:*

ATGACTGTCACAACTTTCTCTTCTACTGGCACAGCCGCCTCCGGGCCTGGCCATGCCCACATCGCCAT
CGTTGGTGCCGGACCTCGCGGGACGAGCGTCGTCGAGCGCCTCGTGGCCTACGCACCCGACCTCATC
ACCTCGGGCAAACAACTCACAGTCCACGTTATCGACCCTTCTTCCCCGGGGCCCGGCAACGTGTGGC
GCACCAACCAGTCCAAAGAGCTCCTGATGAACACCGTCACCTCCCAAATCACATTATTCACCGACAA
GAGCGTCGAGTCGGCGGGTCGCATCGTCCCGGGGCCGACCCTGTACGAGTGGCTGAAGGACTCGCAT
CCGGAGCTCGGGCCCGACGACTACCCGACCCGTGCCCTGTACGGGGAGTACCTGACCTGGGTGTTCA
GCGACGTCGTTTCCCGGGCCCCGCCCAACTTCCACGTCGAGGTCCACGCCACCCGTGCCGTCCGTCT
CGACGACGGCCTCAAGGGCTTCCAGACCCTCGCCCTCGAAGACGGCACGACGCTGACGGGCCTGGC
CGCCGTCGTATTGGCCCAGGGCCACCTGCCGCTCGAGGCGGATCGTGACCAGCGGGCGTTGACCGCC
TACGCGCAGCAGCACCAACTGACCTACATCGCGCCCGTCAACCCGGCCGACGTCGACCTCTCCATCC
TCAAGGCGGACGAGCCCGTCTTCCTGCGCGGCCTCGGCCTGTGCTTCTTCGACTACATGGCCCTCCTG
ACCCTGGGCCGCGGCGGCCGCTTCGAGCGCACCGCACCCAACGGGGGCTTGAGTTATATCGCCTCGG
GTGCCGAGCCCCGGATGTACGCCAGCTCGCGCCGCGGCATCCCCTTCCAGGCGCGCGGCGACAACGA
GAAGGGCGCTTTCGGGCGCCACTACTCCATCCTCATGACGGACGAGGTGATTGCCGACTTCCGGCGC
CGCGCCGGCAAGGGGGACGCGCCCGACTTCATGACCGAGGTCTGGCCCATGATCCGCAAGGAGGTC
GAGCTCATCTACTACGAGGCCCTGCTGCGCCGCGCCGACTTCCGGGACCGGTTCCTGGCCATGGCCTC
GGACAGCGCTGAGGAGACACAGTTGTTGGAGCAGGAGCTCGGGGTGGCGGCGGAGCAGCGCTGGT
CGTGGGAGCGTCTGGGACGACCGCATGCGGCGCATACGTTTACCAACGCCCAGGAGTGGCAGGAGT
GGCTCGTCGGGTACCTGCGCGAGGACGTCAAGCACGCGTCCCTCGGCAATGTCAGCGGGCCCCTGAA
GGCGGCCCTGGATCTGCTGCGCGACCTGCGCAACGAGGTTCGACTTATCGTCGACCACGCCGGCCTC
TCGGGCGAGTCCCGCCGCCAGCACCTTGACCGCTGGTACACCCCCCTGAACGGCTACCTCTCCATTG
GCCCTCCCCGCGAGCGTATCGAGCAGATGATCGCCCTGCTCGAGGCAGGTATCTTGACCCTGGTCGGC
CCCTCCCCCGAAGTGGTCCCGGGTGGATCGGAGCACGAAGATGCCTGGATCGCCCGATCCCCCGAGA
TCCCCGGGTCGGCGGTCCGCGTCACGGCCCTGATCGAGGCTCGTCTGCCTGAGCCGGATCTGCGACA
GACGGGAGATGCGCTGCTGGCGCGGCTACAGGAGACGGGACAGTGCCGTCCTCATGTGGTGGATGG
GTACGAGACGGGCGGACTCGATGTGACGCACAGCCCCTTCCACTTGATCGACTGCCAGGGCCGGGCG
CACCCGCGACGGTTCGCCTTGGGGGTCCCCACTGAGGGGGCGCATTGGGTCACCGCGGCGGGGGCC
CGGCCGGGGGTGAACTCGGTCACGCTTTGCGATACTGATGCTGTGGCCGGGGCCGCGCTCGTGACTG
CATCCAGTGGCCTCAGGGGTCTCTCGCCGGAGTTCCGGCCGATGGTGCAGACCGAGTCGATGATGAT
CCAGCCGATCCGTCTATAA

**Protein sequence of NpaA homolog as-NpaA.**

as-NpaA:

MTVTTFSSTGTAASGPGHAHIAIVGAGPRGTSVVERLVAYAPDLITSGKQLTVHVIDPSSPGPGNVWRTNQ
SKELLMNTVTSQITLFTDKSVESAGRIVPGPTLYEWLKDSHPELGPDDYPTRALYGEYLTWVFSDVVSRA
PPNFHVEVHATRAVRLDDGLKGFQTLALEDGTTLTGLAAVVLAQGHLPLEADRDQRALTAYAQQHQLTY
IAPVNPADVDLSILKADEPVFLRGLGLCFFDYMALLTLGRGGRFERTAPNGGLSYIASGAEPRMYASSRRG

IPFQARGDNEKGAFGRHYSILMTDEVIADFRRRAGKGDAPDFMTEVWPMIRKEVELIYYEALLRRADFR
DRFLAMASDSAEETQLLEQELGVAAEQRWSWERLGRPHAAHTFTNAQEWQEWLVGYLREDVKHASLG
NVSGPLKAALDLLRDLRNEVRLIVDHAGLSGESRRQHLDRWYTPLNGYLSIGPPRERIEQMIALLEAGILT
LVGPSPEVVPGGSEHEDAWIARSPEIPGSAVRVTALIEARLPEPDLRQTGDALLARLQETGQCRPHVVDGY
ETGGLDVTHSPFHLIDCQGRAHPRRFALGVPTEGAHWVTAAGARPGVNSVTLCDTDAVAGAALVTASSG
LRGLSPEFRPMVQTESMMIQPIRL

**DNA sequence of *npaB* homolog *as-npaB* gene.**

*as-npaB:*

ATGTCGACCAATCCCAAGTTTGACGAGCTGCACAAGCAGCTGTTCGACGAGGGCGTCAAGGTTCGCC
GCGCTGTCCTCGGAGATGAATATGTCGACAAGGCCCTGCAGAATGCCACCCCTTTCACGATGCCAGGC
CAGCAACTCATTACTGAGTGGGCCTGGGGGACCGTCTGGCAGCGTCCAGGCCTGGACCGGAAGCAG
CGCAGTCTCCTGAACATCGGCATCATCATTGCCCAGAAGGCGTGGCTCGAACTGGAGCTCCATACCCG
CGGCGCCATCAACAATGGGCTGACCGAGGTCGAGATCCGCGAGGCAGTTCTGCAAGCCACCGTCTAT
TGCGGCACACCCGCCGGCGTCGAGGCCATGATGGTGACGGAAAAGACCATCAACGAGATGATTGCCA
AGGGGGAGTACACAAGGCCGGCTTGA

**Protein sequence of NpaB homolog as-NpaB.**

as-NpaB:

MSTNPKFDELHKQLFDEGVKVRRAVLGDEYVDKALQNATPFTMPGQQLITEWAWGTVWQRPGLDRKQ
RSLLNIGIIIAQKAWLELELHTRGAINNGLTEVEIREAVLQATVYCGTPAGVEAMMVTEKTINEMIAKGEY
TRPA

**DNA sequence of synthesized *mr-npaA* gene.**

*mr-npaA*:

ATGACGGCGCTGTTGCGTAATGAATGTAAGCAGGCCTTGGCAAAAAGCTTACCCTGGACCAAATCAC
CCCTGGTGATTTCTGCTCCTATGCGTGTCATGACCGGCCCTGGTCTTGCCGTGGCTGTAAGTAGTGCG
GGGGGATTGGGCTTCCTGGGTCCCACGCTGAAGCCTGAAGATGTCTTCGCTGACCTTGACAAGGCTG
CAGAGCTTCTTGGTAGTTCGCCGATTCAAGGTGCTGCGGGACCGTCCTTGCTTCCAATCGGCGTGGGC
TTTCAGACATGGAACGGCGATTTGGAGGTTGCGGTATCCGCGGTGACTAAGCACCGCCCCTGCGCGG
TGTGGCTGTTCGCGCCGCGCCGCGGGCAAGCAGAGTTCAATGAGTGGACAGCGGCTATTCGCTCAGC
TAGTCCGGACACGCGCGTATGGCTTCAGGTGGGGAGCTTGGGAGAAGCGGTGGAAGCAGCAGCATC
AGCCACCCCACCCGACGTCCTGGTGCTTCAAGGAGCGGAAGGAGGCGGACATGGTCGTCACCGCGA
CGCGCAGGGCACCATCGCTCTTGTACCCGAGGTTTCGGACGCACTTGGTCACTCTGGGATCCCGCTG
GTTGCAGCCGGGGGGATTGTTGATGGCCGTGGGGCTGCGGCGGCTTTGACCTTAGGTGCGGCTGGGG
TCGCGATGGGTACACGTTTTCTTGCTTCATCAGAGGCTCGCATCTCTAAAGGTTATCAAGACGAGGTT
GTACGTGCATCAGATGGAGCAAAGAACACCGTCCGCACCCAATTATACAATCACCTGCGCGGTACGTT
TGGTTGGCCCGAACCGTTCTCTCCTCGCACCCTTATTAATCGCTCGTGGCGCGATCACGAAGCTGGCG
TTGAGTTCGACCGTTTGAAGGAGTTACACGATGAGTCGGCTAAGACGGGTGACGCCGGATGGGGGCC
TGAGGGACGCTTAGCTACATACGTCGGCGCGGCGGTGGGCTTAGTCCGCCGCGTCGACGACGCTGCA
GTGATTGTTCGTGAGACCCGCGATCAGGCACGCGCCATTCTTACTTCGGTAGTCGCACATTTA

#### Supplementary References


(1) Jin, F. J.; Maruyama, J.; Juvvadi, P. R.; Arioka, M.; Kitamoto, K. Development of a Novel Quadruple Auxotrophic Host Transformation System by argB Gene Disruption Using adeA Gene and Exploiting Adenine Auxotrophy in Aspergillus Oryzae. *FEMS Microbiol. Lett.* **2004**, *239* (1), 79–85.

(2) Liu, N.; Hung, Y.-S.; Gao, S.-S.; Hang, L.; Zou, Y.; Chooi, Y.-H.; Tang, Y. Identification and Heterologous Production of a Benzoyl-Primed Tricarboxylic Acid Polyketide Intermediate from the Zaragozic Acid A Biosynthetic Pathway. *Org. Lett.* **2017**, *19* (13), 3560–3563.

(3) 精二中村; 忠次郎下田. 麹菌の生産する抗生物質 Oryzacidin に関する研究. 日本農芸化学会誌 **1954**, *28* (11), 909–913.

(4) Turgeon, B. G.; Condon, B.; Liu, J.; Zhang, N. Protoplast Transformation of Filamentous Fungi. In *Molecular and Cell Biology Methods for Fungi*; Sharon, A., Ed.; Methods in Molecular Biology; Humana Press: Totowa, NJ, 2010; pp 3–19.

(5) Ohashi, M.; Liu, F.; Hai, Y.; Chen, M.; Tang, M.; Yang, Z.; Sato, M.; Watanabe, K.; Houk, K. N.; Tang, Y. SAM-Dependent Enzyme-Catalysed Pericyclic Reactions in Natural Product Biosynthesis. *Nature* **2017**, *549* (7673), 502–506.

(6) Sugai, Y.; Katsuyama, Y.; Ohnishi, Y. A Nitrous Acid Biosynthetic Pathway for Diazo Group Formation in Bacteria. *Nat. Chem. Biol.* **2016**, *12* (2), 73–75.

(7) Gilchrist, C. L. M.; Booth, T. J.; van Wersch, B.; van Grieken, L.; Medema, M. H.; Chooi, Y.-H. Cblaster: A Remote Search Tool for Rapid Identification and Visualization of Homologous Gene Clusters. *Bioinform. Adv.* **2021**, *1* (1), vbab016.

(8) Notredame, C.; Higgins, D. G.; Heringa, J. T-Coffee: A Novel Method for Fast and Accurate Multiple Sequence Alignment1. *J. Mol. Biol.* **2000**, *302* (1), 205–217.

(9) Waterhouse, A. M.; Procter, J. B.; Martin, D. M. A.; Clamp, M.; Barton, G. J. Jalview Version 2—a Multiple Sequence Alignment Editor and Analysis Workbench. *Bioinformatics* **2009**, *25* (9), 1189–1191.

(10) Valentino, H.; Sobrado, P. Characterization of a Nitro-Forming Enzyme Involved in Fosfazinomycin Biosynthesis. *Biochemistry* **2021**, *60* (38), 2851–2864.

(11) Gadda, G.; Francis, K. Nitronate Monooxygenase, a Model for Anionic Flavin Semiquinone Intermediates in Oxidative Catalysis. *Arch. Biochem. Biophys.* **2010**, *493* (1), 53–61.

(12) Salvi, F.; Agniswamy, J.; Yuan, H.; Vercammen, K.; Pelicaen, R.; Cornelis, P.; Spain, J. C.; Weber, I. T.; Gadda, G. The Combined Structural and Kinetic Characterization of a Bacterial Nitronate Monooxygenase from Pseudomonas Aeruginosa PAO1 Establishes NMO Class I and II *. *J. Biol. Chem.* **2014**, *289* (34), 23764–23775.
